# Male germ cell-associated kinase (MAK) is required for axoneme formation during ciliogenesis in zebrafish photoreceptors

**DOI:** 10.1101/2023.11.19.567778

**Authors:** Hung-Ju Chiang, Yuko Nishiwaki, Wei-Chieh Chiang, Ichiro Masai

**Affiliations:** Developmental Neurobiology Unit, Okinawa Institute of Science and Technology Graduate University, Tancha, Okinawa, Japan

**Keywords:** zebrafish, photoreceptor degeneration, MAK, ciliopathy, ciliogenesis

## Abstract

Vertebrate photoreceptors are highly specialized retinal neurons that have cilium-derived membrane organelles called outer segments (OS), which function as platforms for phototransduction. Male germ cell-associated kinase (MAK) is a cilium-associated serine/threonine kinase, and its genetic mutation causes photoreceptor degeneration in mice and retinitis pigmentosa in humans. However, the role of MAK in photoreceptors is not fully understood. Here, we report that zebrafish *mak* mutants show rapid photoreceptor degeneration during embryonic development. In *mak* mutants, both cone and rod photoreceptors completely lack OSs and undergo apoptosis. Interestingly, zebrafish *mak* mutants fail to generate axonemes during photoreceptor ciliogenesis, whereas basal bodies are specified. These data suggest that MAK contributes to axoneme development in zebrafish, in contrast to mouse *Mak* mutants, which have elongated photoreceptor axonemes. Furthermore, the kinase activity of MAK is critical in ciliary axoneme development and photoreceptor survival. Thus, MAK is required for ciliogenesis and OS formation in zebrafish photoreceptors to ensure intracellular protein transport and photoreceptor survival.

**Summary statement:** Male germ cell-associated kinase (MAK) is a cilium-associated serine/threonine kinase that promotes axoneme development during ciliogenesis in zebrafish photoreceptors to ensure intracellular protein transport and photoreceptor survival.

## Introduction

The vertebrate photoreceptor is highly compartmentalized to form specialized structures related to phototransduction (Fain et al., 2010). The outer segment (OS) is a specialized cilium of the photoreceptor, in which multiple photoreceptive membrane discs are regularly stacked to accommodate phototransduction molecules such as rhodopsin and opsins (Bachmann-Gagescu and Neuhauss, 2019; Chen et al., 2021). The inner segment (IS) is a mitochondria-enriched region between the OS and the nucleus. Finally, photoreceptors have a specialized synaptic structure in the most basal region that mediates transmission of electric signals to horizontal cells and bipolar cells (Mercer and Thoreson, 2011).

The photoreceptor OS is a highly specialized primary cilium that consists of stacked membrane discs around a microtubule-based backbone called the axoneme (Bachmann-Gagescu and Neuhauss, 2019; Chen et al., 2021). The axoneme is anchored to the basal body in the IS of photoreceptors and extends apically through the connecting cilium. The connecting cilium bridges the IS and the OS, through which phototransduction molecules are transported from the ER to the Golgi and then into the OS. Importantly, the connecting cilium is equivalent to the ciliary transition zone in other cell types and functions as a gating system (Park and Leroux, 2022), through which OS resident proteins are transported to the OS by the intraflagellar transport (IFT) complex (Klena and Pigino, 2022) and BBSome, composed of eight Bardet-Biedl syndrome (BBS) proteins (Tian et al., 2023).

The process of ciliary development is called ciliogenesis, which consists primarily of three steps: centriole (basal body) docking to the apical plasma membrane, establishment of a transition zone, and ciliary axoneme extension (Breslow and Holland, 2019; Chen et al., 2021). The first step of ciliogenesis is removal of CP110 from a distal appendage of the mother centriole, which enables ARL13b and components of IFT-A and IFT-B to assemble at the apical surface of a distal appendage in a rab8-dependent manner (Breslow and Holland, 2019; Ge et al., 2022). The second step is formation of the transition zone by transport of MKS and NPHP module components (Park and Leroux, 2022). The third step is extension of the axoneme through IFT-mediated ciliary transport, which transports and adds α-/β-tubulin dimers to the plus end of the microtubule comprising the axoneme (Brouhard and Rice, 2018; Ge et al., 2022; Nakayama and Katoh, 2018). Defects in ciliogenesis cause ciliopathy, which is associated with various abnormalities during organogenesis, including retinal dystrophies, cystic kidney diseases, skeletal dysplasia, and polydactyly (Mill et al., 2023). Although many factors are reportedly involved in photoreceptor ciliogenesis and ciliopathy (Cideciyan et al., 2007; den Hollander et al., 2006; Gao et al., 2002; Gorden et al., 2008; Kakakhel et al., 2020; Rao et al., 2015; Zhu et al., 2021), the underlying mechanisms are not fully understood.

Male germ cell-associated kinase (MAK) belongs to the MAK/ICK/MOK serine/threonine kinase family (Chen et al., 2013). MAK was identified by cross-hybridization with tyrosine kinase v-ros in rat testicular cells, so it was considered to be a spermatogenesis regulator (Matsushime et al., 1990). MAK functions as a coactivator of androgen receptors (ARs) to promote AR-mediated signaling, which is associated with prostate tumorigenesis (Ma et al., 2006; Wang and Kung, 2012; Xia et al., 2002). Furthermore, *Mak* transcripts are expressed in the photoreceptor layer in mouse retina (Blackshaw et al., 2004) and MAK protein is localized in cilia of photoreceptors (Omori et al., 2010). In *Mak* knockout mice, photoreceptors show abnormally elongated axonemes and malformation of membrane discs in the OS, leading to photoreceptor degeneration (Omori et al., 2010). In humans, *MAK* mutations cause retinitis pigmentosa (RP), in which patients progressively lose their vision (Ozgul et al., 2011; Stone et al., 2011). However, human patients carrying *MAK* mutations do not show elongation of the photoreceptor layer (Stone et al., 2011), although fibroblasts derived from RP patients carrying *MAK* mutations have elongated cilia (Tucker et al., 2022). Thus, it is important how *mak* mutations affect ciliary regulation in photoreceptors among vertebrate species.

In this study, we found that zebrafish photoreceptors of *mak* mutants degenerate during embryonic development. In *mak* mutants, both rods and cones undergo apoptosis, although their degeneration processes differ. Interestingly, *mak* mutants fail to form axonemes in photoreceptor cilia, whereas basal bodies are specified, which is the opposite of mouse *Mak* knockout photoreceptors, with elongated ciliary axonemes. Furthermore, both cones and rods completely lack OSs in *mak* mutants, leading to ectopic distribution of opsins. So, MAK is essential to form axonemes and OSs in photoreceptors. Finally, MAK kinase activity is critical for axoneme formation and photoreceptor survival. Thus, MAK is essential for ciliogenesis, OS formation, and photoreceptor survival.

## Results

### *Z*ebrafish *pday* mutants carry a mutation in *mak* gene, leading to photoreceptor degeneration

Zebrafish *payday* (*pday*) mutants display no visual response during the embryonic stage (Muto et al., 2005). We selected *pday* homozygous mutant embryos using an optokinetic response (OKR) assay and examined their phenotypes. At 6 days post-fertilization (dpf), *pday* mutants showed no apparent morphological defect (Fig. 1A), but did not survive beyond 10 dpf, suggesting a lethal mutation. However, under intensive feeding, less than 1% of *pday* mutants survived until 2 months post-fertilization (mpf), displaying scoliosis, cardiac edema, and smaller body size (Fig. 1A).

**Fig. 1:**
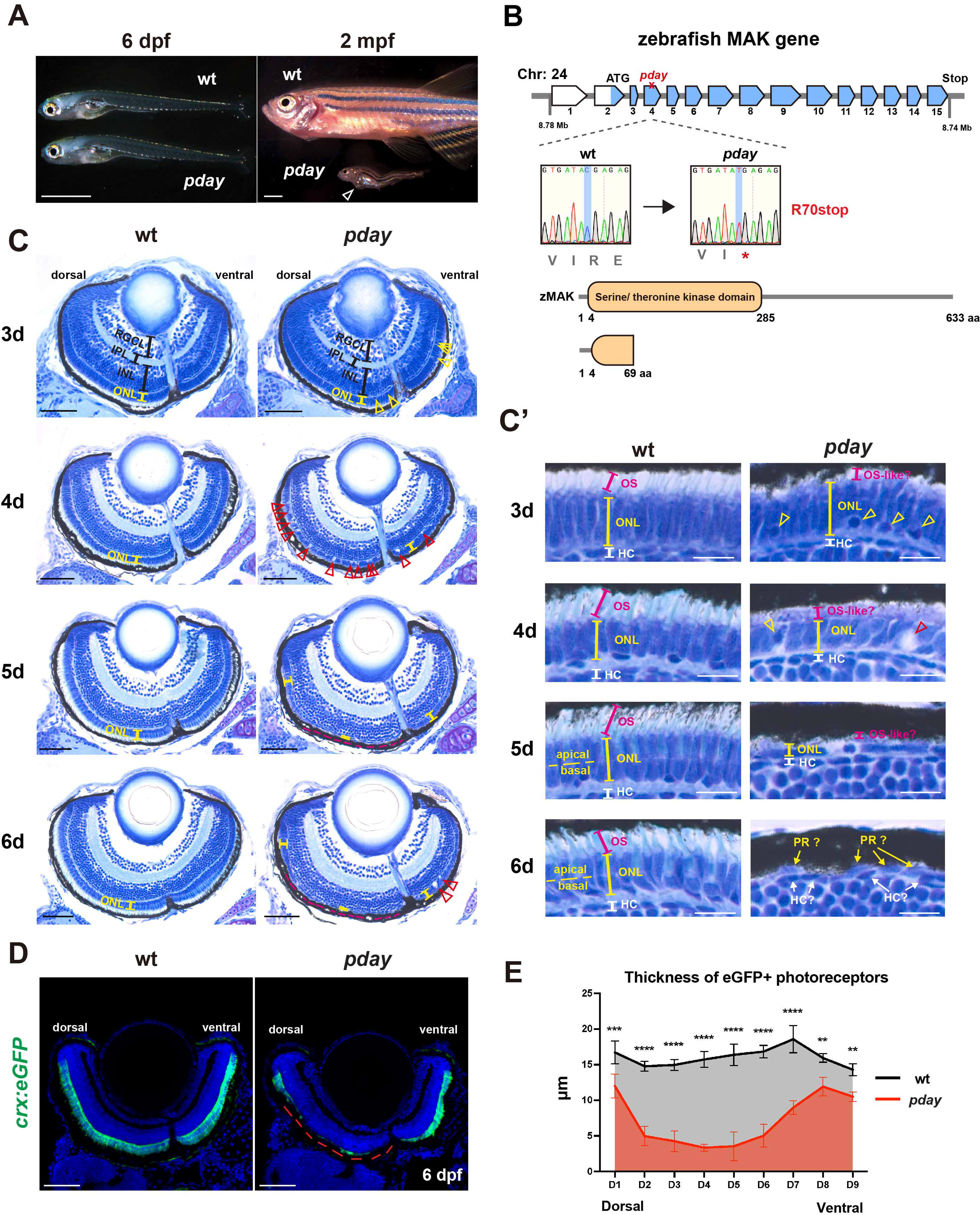
Zebrafish *mak* mutants, *pday,* show photoreceptor degeneration. (A) *pday* mutants at 6 dpf and 2 mpf. At 6 dpf, *pday* mutants show no morphological differences from wild-type siblings. At 2 mpf, *pday* mutants have smaller bodies with less-developed fins, scoliosis, and cardiac edema (open arrowhead). (B) The zebrafish *mak* gene consists of 15 exons and encodes 633 amino acids with a serine/threonine kinase domain. In *pday* mutants, a non-sense mutation (R70stop) results in a truncated protein lacking most of kinase domain. (C) Wild-type and *pday* mutant retinas. The ONL in *pday* mutants shows dense nuclear granules (yellow arrowheads), bubble-like structures (red arrowheads) and reduced outer segments (yellow lines). In *pday* mutants, the dorso-central ONL progressively decreases in thickness after 4 dpf, becomes extremely flat at 5 dpf and disappears at 6 dpf (red dotted lines). (C’) Higher-magnification of the central ONL. The OS is drastically reduced in *pday* mutants (red lines). Dense nuclear granules (yellow arrowheads), bubble-like structures (red arrowheads), and the progressive reduction of ONL thickness (yellow lines) are observed in *pday* mutants. HC, horizontal cells; PR, photoreceptors. (D) Wild-type and *pday* mutant retinas carrying the transgenic line *Tg[crx:eGFP]*. Nuclei were counterstained with Hoechst 33342 (blue). crx:eGFP (green) expression is more severely decreased in the dorsal retina (red dotted line) than in the ventral retina of *pday* mutants. (E) Thickness of eGFP-positive photoreceptors along the DV axis of the retina. The thickness is more severely decreased in the dorsal retina than in the ventral retina of *pday* mutants. Color bars and lines indicate mean ± SD. Statistical difference is evaluated with 2-way ANOVA and Sidak’s multiple comparison tests, n=3 for each point. ** p<0.01, *** p<0.001, **** p<0.0001. Scale bars: 1 mm (A), 50 µm (C), 10 µm (C’), 40 µm (D).

Next, we mapped the *pday* mutant locus with simple sequence-length polymorphism (SSLP) markers (Knapik et al., 1998; Shimoda et al., 1999). The *pday* mutation was mapped between two SSLP markers z23011 (1/186 miosis) and z13695 (3/186 miosis), so the mutant gene is restricted to a genomic region between 8.517 and 9.123 Mb on chromosome 24. In this genomic region, 10 genes including *mak* were annotated in the zebrafish genomic database (GRCz10, Ensemble release 80). Since *Mak* mutations cause photoreceptor degeneration in mice (Omori et al., 2010) and humans (Ozgul et al., 2011; Stone et al., 2011), and because the other 9 genes were not reported to be involved in retinal development, degeneration, and functions, we focused on the *mak* gene. First, we designed a new polymorphic marker, called Mak-N1, located around 20 bp upstream of the first exon of the *mak* gene. The recombination rate of Mak-N1 for *pday* mutation was 0 of 186 meiosis, suggesting that *mak* is a strong candidate. Next, we cloned *mak* cDNA prepared from wild-type embryos, and found that *mak* cDNA comprises 15 exons (Fig. 1B) and that there are two isoforms of *mak* mRNA. One has and the other lacks exon 14 due to alternative splicing. Then, we focused on the longest isoform, which has exon 14 and is annotated as the full-length isoform in the zebrafish genomic database (mak-201, GRCz11, Ensemble release 110). We compared amino acid sequences of this longest zebrafish MAK isoform with those of human and mouse MAK. Amino acid identity between zebrafish MAK and human/mouse MAK was more than 50% for full-length protein and more than 86% for the N-terminal kinase domain (Fig. S1). We cloned *mak* cDNAs from *pday* mutant mRNA and genomic fragments covering coding exons from the *pday* mutant genome. Sequencing revealed that a non-sense mutation occurs in exon 4 of the *mak* gene in *pday* mutants, which causes a premature termination codon at 70R (Fig. 1B). These data suggest that the *pday* mutant gene encodes MAK.

Next, we examined retinal phenotypes of wild-type and *pday* mutant embryos from 3 to 6 dpf (Fig. 1C, 1C’). In wild-type retinas at 3 dpf, photoreceptors differentiate to form the outer nuclear layer (ONL). The OS is formed in the apical ONL (Fig. 1C’). In *pday* mutants, the ONL was formed; however, densely labeled nuclei were observed in the basal region of the ventro-central ONL (Fig. 1C, 1C’), and the OS was markedly reduced (Fig. 1C’). In wild-type photoreceptors at 4 dpf, the OS was elongated. However, in *pday* mutants, the ONL was decreased in thickness (Fig. 1C, 1C’), and no OS-like structure was observed (Fig. 1C’). In addition, there were pyknotic nuclei and bubble-like structures (Fig. 1C, 1C’). In wild-type embryos at 5 dpf, the ONL was divided into two subnuclear layers (Fig. 1C’), the apical and basal regions of which contain nuclei of blue/red/green cones and rods/UV cones, respectively (D’Orazi et al., 2020; Oel et al., 2020). However, in *pday* mutants, the dorso-central ONL drastically decreased in thickness (Fig. 1C) and contained flattened nuclei (Fig. 1C, 1C’). In wild-type embryos at 6 dpf, photoreceptors showed more mature shapes in the ONL. However, in *pday* mutants, most photoreceptor nuclei disappeared in the dorso-central retina (Fig. 1C), so there are very flat nuclei in the outermost region of the retina, which are difficult to define as the ONL or horizontal cells (Fig. 1C’). Interestingly, in *pday* mutants, ONL thickness was less affected in the retinal region more ventral to the optic nerve head, even at 6 dpf (Fig. 1C). Thus, sensitivity of photoreceptor degeneration differs between the dorso-central and the ventral retina. In addition, the inner nuclear layer (INL), the inner plexiform layer (IPL), and the retinal ganglion cell layer (RGCL) seemed to be intact in *pday* mutants from 3 to 6 dpf (Fig. 1C), suggesting photoreceptor-specific degeneration in *pday* mutants.

To confirm photoreceptor degeneration in mutant retinas at 6 dpf, we introduced a transgenic line *Tg[crx:eGFP]* that expresses eGFP in photoreceptor precursors and mature rod and cone photoreceptors under control of the *cone-rod homeobox* (*crx*) promoter (Suzuki et al., 2013) (Fig. 1D). In wild-type retinas, eGFP signals were detected in the ONL at 6 dpf. However, in *pday* mutant retinas, eGFP signals disappeared in the dorso-central retina (Fig. 1D). We measured thicknesses of eGFP-positive areas along the dorso-ventral axis (Fig. S2A), and found that photoreceptor degeneration is more severe in the dorso-central retina than in the ventral retina (Fig. 1E). Thus, photoreceptors undergo degeneration in *pday* mutants.

### Both rods and cones degenerate in *pday* mutants

In human *MAK*-associated RP patients, rods degenerate more prominently (Ozgul et al., 2011; Stone et al., 2011). To evaluate rod and cone degeneration separately, we generated double transgenic lines, *Tg[rho:NLS-eGFP; gnat2:NLS-tdTomato*], to visualize rods and cones with eGFP and tdTomato fluorescence, respectively, and introduced them into *pday* mutants (Fig. 2, Fig. S3A). In wild-type retinas, eGFP expression in rods was strong and dense in the ventral ONL but relatively weak and sparse in the dorso-central ONL at 3 dpf (Fig. S3A). On the other hand, eGFP expression was very faint in *pday* mutant retinas under the same conditions. However, when laser intensity was increased, eGFP expression was detected (Fig. S3A), suggesting that rods are maintained in 3 dpf *pday* mutant retinas, although the rhodopsin promoter does not effectively drive *eGFP* mRNA or else translation of *eGFP* mRNA to eGFP protein was affected in *pday* mutants. Therefore, in later experiments using *Tg[rho:NLS-eGFP]*, we applied higher laser intensity to *pday* mutant scanning, and compared rod phenotypes between *pday* mutants and wild-type siblings.

**Fig. 2:**
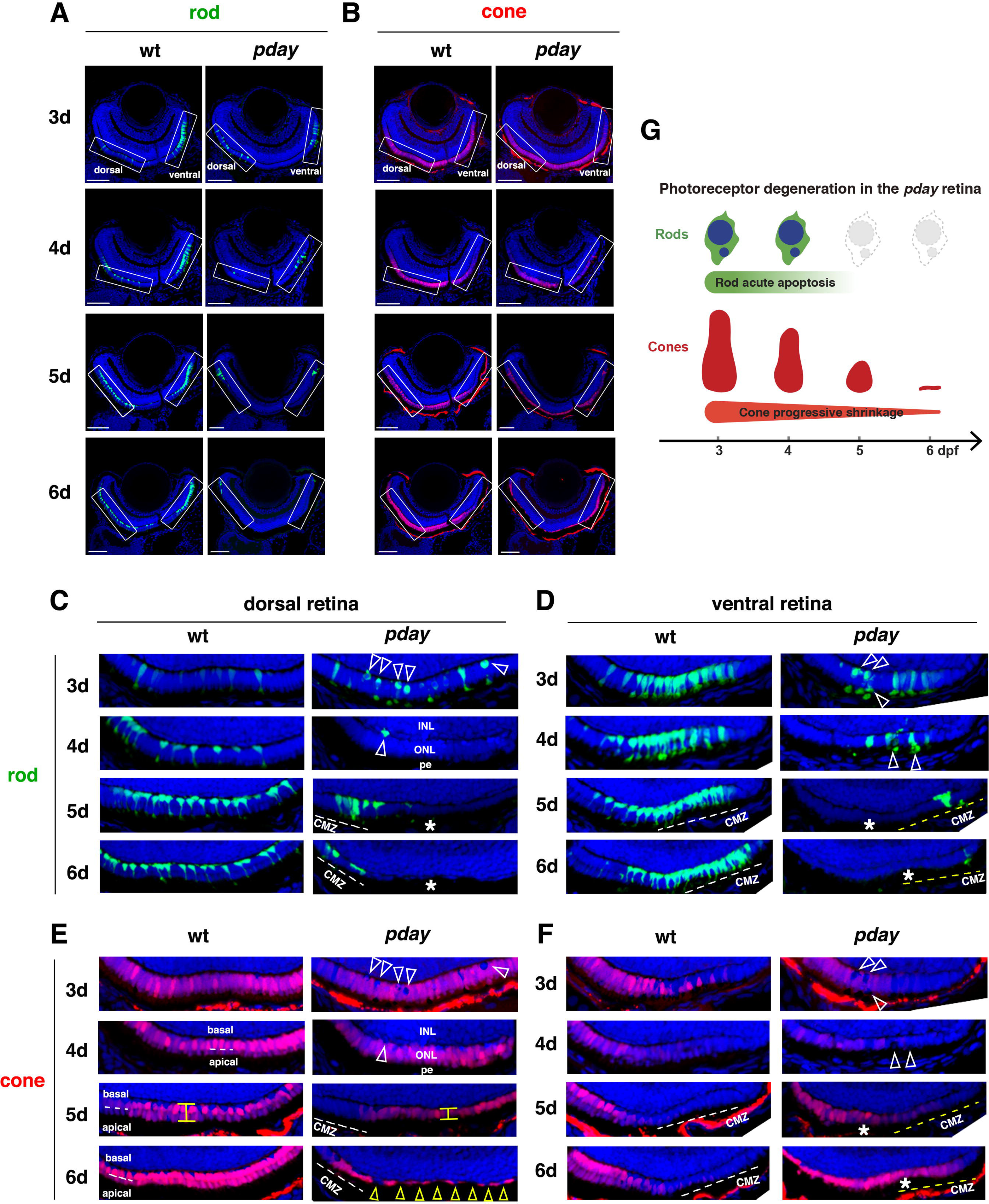
Rods undergo apoptotic-like degeneration in *pday* mutants, whereas cones progressively shrink. (A-B) Wild-type and *pday* mutant retinas carrying the transgenic lines *Tg[rho:NLS-eGFP]* and *Tg[gnat2:NLS-tdTomato]*, which visualize rods (green) and cones (red), respectively. Nuclei were counterstained with Hoechst 33342 (blue). (A) Only green and blue channel. (B) Only red and blue channel. Squares indicate the dorsal and ventral ONL, shown in (C–F). (C) The dorsal ONL of wild-type and *pday* mutant retinas with *Tg[rho:NLS-eGFP]* (green). In *pday* mutants, rods have condensed, round nuclei (white arrowheads) at 3 dpf. In *pday* mutants, rods are drastically reduced in number at 4 dpf and disappear at 5 and 6 dpf (asterisks), except in the peripheral region near the dorsal CMZ. (D) The ventral ONL of wild-type and *pday* mutant retinas with *Tg[rho:NLS-eGFP]* (green). In *pday* mutants, rods have condensed, round nuclei (white arrowheads) at 3 and 4 dpf. In *pday* mutants, rods disappear at 5 and 6 dpf (asterisks), except in the peripheral region near the ventral CMZ. (E) The dorsal ONL of wild-type and *pday* mutant retinas with *Tg[gnat2:NLS-tdTomato]* (red). In wild type, cones form monolayer at 3 dpf and two subnuclear layers after 4 dpf. In *pday* mutants, tdTomato-negative nuclei are observed, whose position is identical to that of condensed rod nuclei (white arrowheads). From 4 to 5 dpf, cones shrink progressively and become flattened (yellow line). At 6 dpf, cones are extremely flattened and disappear, leaving gaps (yellow arrowheads), except in the peripheral region near the dorsal CMZ. (F) The ventral ONL of wild-type and *pday* mutant retinas with *Tg[gnat2:NLS-tdTomato]* (red). In wild type, cones differentiate at 3 dpf, but progressively disappear toward the periphery of the ventral ONL, where rods densely differentiate. In *pday* mutants, ONL thickness is progressively decreased after 4 dpf. Interestingly, after 5 dpf, the cone differentiating area is expanded toward the periphery of the ventral ONL (asterisks), where rods disappear. (G) Cone and rod degeneration process in *pday* mutants. Rods undergo acute degeneration with apoptotic-like pyknotic nuclei during 3-4 dpf, whereas cones undergo progressive shrinkage of cell volume during 4-6 dpf and disappear by 6 dpf. Scale bars: 40 µm (A, B).

Since photoreceptor degeneration is more severe in the dorso-central retina than the ventral retina at 6 dpf (Fig. 1D, 1E), we visualized morphology of rods and cones using *Tg[rho:NLS-eGFP; gnat2:NLS-tdTomato*] from 3 to 6 dpf (Fig. 2A, 2B), and focused on the dorsal and ventral ONL regions (Fig. 2C-F). First, we examined rods (Fig. 2A). In wild-type retinas at 3 dpf, rods were sparse in the dorsal ONL (Fig. 2C), but densely produced in the ventral ONL (Fig. 2D). In *pday* mutants, many rods show very round nuclei in both the dorsal and ventral ONL (Fig. 2C and 2D). Interestingly, eGFP signals were predominantly localized in nuclei of wild-type rods because eGFP was tagged with a nuclear localization sequence (NLS); however, eGFP signals were often excluded from the round nuclei of *pday* mutant rods, suggesting apoptotic-like pyknotic nuclei of *pday* mutant rods. After 4 dpf, in wild-type retinas, rods progressively increased in number and aligned along the dorsal ONL. Their nuclei are positioned in the most basal region of the ONL, and rods extend processes to form the OS in the apical ONL (Fig. 2C). On the other hand, rods are more densely differentiated in the ventral ONL (Fig. 2D). However, in *pday* mutants at 4 dpf, rods almost disappeared in both the dorsal and ventral ONL. Only a few rods remained, but displayed pyknotic nuclei or were excluded from the ONL (Fig. 2C and 2D). In *pday* mutants at 5 and 6 dpf, rods disappeared in the ONL (Fig. 2C and 2D), except in the peripheral retina adjacent to the ciliary marginal zone (CMZ) where new photoreceptors continued to be produced by retinal stem cells. So, rods are eliminated soon after they are specified from the CMZ. Evaluation of numbers of rod nuclei along the DV axis of the retina confirmed that rods were progressively diminished in the central ONL of *pday* mutant retinas and were completely eliminated, except in the CMZ area at 6 dpf (Fig. S3B).

Next, we examined cones (Fig. 2B). In wild-type retinas at 3 dpf, cones differentiated uniformly in the dorsal ONL (Fig. 2E); however, cone density became progressively sparse toward the CMZ region in the ventral ONL (Fig. 2F), a pattern complementary to that of rods (Fig. 2D). In *pday* mutant retinas, cones normally differentiated to form a spatial pattern similar to that of wild-type retinas (Fig. 2E, 2F). However, there were tdTomato-negative rounded nuclei in both the dorsal and ventral ONL (Fig. 2E and 2F), whose positions are identical to pyknotic-like rod nuclei (Fig. 2C and 2D). Thus, rods lose their normal localization in the basal ONL in *pday* mutants at 3 dpf. After 4 dpf, in wild-type retinas, cones form two subnuclear layers in the dorso-central ONL (Fig. 2E). However, cone density was complementary to rod density and became progressively sparse toward the CMZ region in the ventral ONL (Fig. 2F). In *pday* mutants at 4 dpf, segregation of two cone subnuclear layers was incomplete in the dorso-central ONL (Fig. 2E). Furthermore, cones were sparse and shortened in the ventral ONL (Fig. 2F). In *pday* mutants at 5 dpf, cones were shortened in the dorsal ONL (Fig. 2E). Interestingly, cone density increased toward the CMZ in the ventral ONL of *pday* mutants (Fig. 2F), where rods failed to be maintained (Fig. 2D). In *pday* mutants at 6 dpf, cones were extremely flattened and there were gaps between them in the dorsal ONL (Fig. 2E), indicating that cones had been eliminated. Furthermore, cones were similarly flattened in the ventro-central ONL (Fig. 2F), whereas cones were uniformly produced and less flattened toward the CMZ of the ventral ONL (Fig. 2F). Therefore, rods undergo acute apoptosis-like cell death in *pday* mutants from 3 to 4 dpf, whereas cones undergo progressive shrinkage after 4 dpf and eventually disappear by 6 dpf in *pday* mutants (Fig. 2G), although cone degeneration is less severe in the peripheral region of the ventral ONL.

### Both rods and cones undergo apoptosis in *pday* mutants

Condensed rod nuclei in *pday* mutants at 3 dpf (Fig. 2C, 2D) are reminiscent of apoptosis, in which chromatin condensation is a hallmark (Oberhammer et al., 1994). We performed TdT-mediated dUTP-nick-end labeling (TUNEL) to *pday* mutant retinas at 3, 4, and 5 dpf (Fig. 3A, 3A’, Fig. S4A). In wild-type retinas, the number of TUNEL-positive cells was continuously very low in the ONL (Fig. 3A, Fig. S4A). In *pday* mutant retinas at 3 dpf, TUNEL signals were detected in the central and ventral ONL (Fig. 3A’), which is consistent with our observation that rod pyknotic nuclei were detected at 3 dpf (Fig. 2C, 2D), suggesting that these early TUNEL-positive cells are likely to be rods. In *pday* mutant retinas at 4 and 5 dpf, TUNEL signals increased in the dorsal and central ONL (Fig. 3A’). Interestingly, TUNEL-positive cells were positioned in a very thin central ONL of 5 dpf *pday* mutant retinas (Fig. 3A’), suggesting that these late TUNEL-positive cells are likely to be cones. TUNEL-positive cells were more numerous in *pday* mutant ONL than wild-type ONL at 3 dpf, but the difference was not significant. However, the number of TUNEL-positive cells was the highest in *pday* mutant ONL at 4 dpf and was still significantly higher in *pday* mutant ONL at 5 dpf, compared with wild-type ONL (Fig. S4A). The number of TUNEL signals in other retinal layers, INL, and RGCL, was not significantly different between *pday* mutants and wild-type siblings at stages 3, 4, or 5 dpf (Fig. S4B, S4C), suggesting that apoptosis is specific to photoreceptors.

**Fig. 3:**
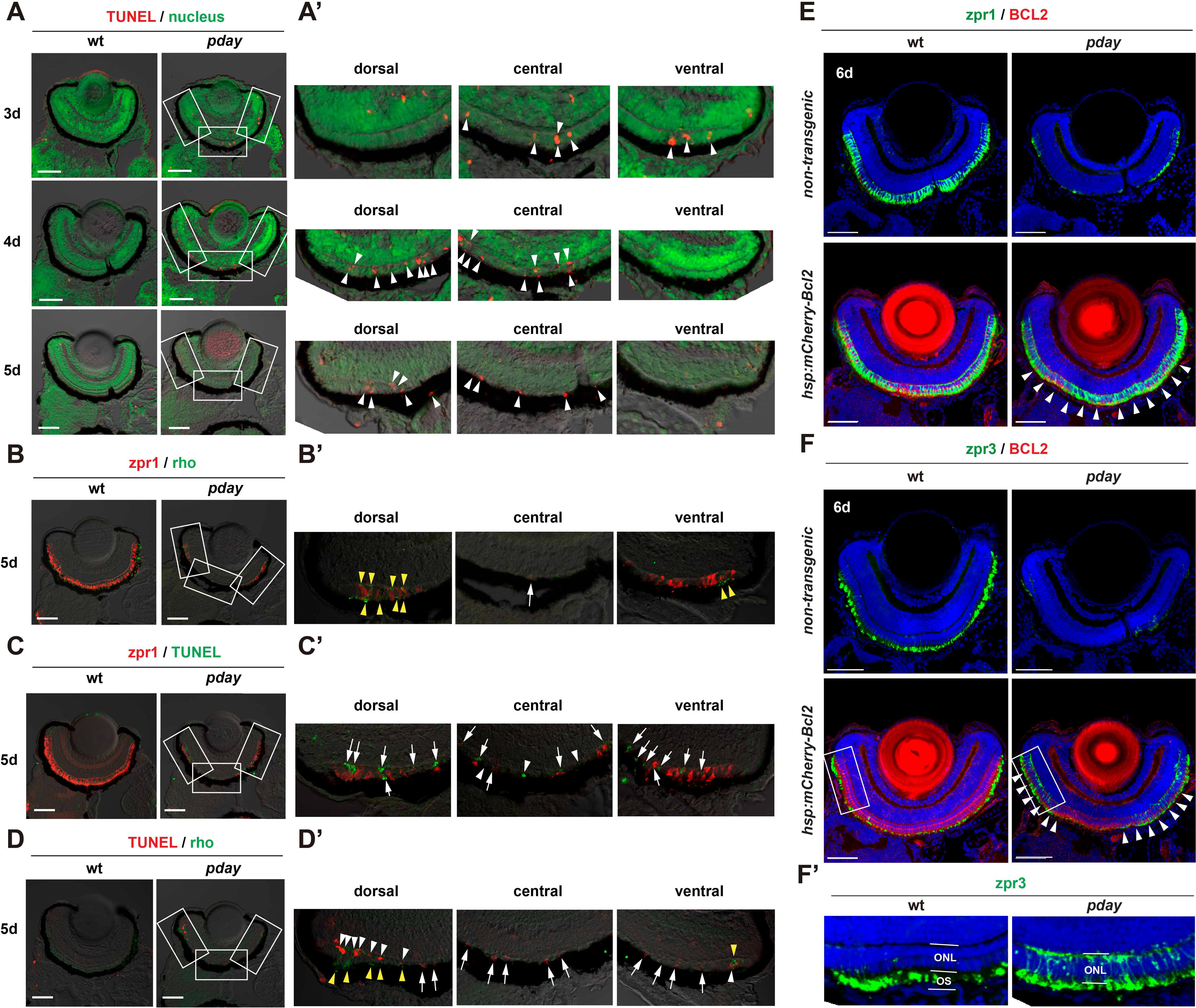
Rods and cones undergo apoptosis in *pday* mutants. (A) TUNEL (red) of wild-type and *pday* mutant retinas. Nuclei are counterstained with Sytox-Green (green). (A’) Higher-magnification of dorsal, central, and ventral ONL of *pday* mutant retinas, which are indicated by white squares in (A). White arrowheads indicate TUNEL signals (red). (B) Double labeling of 5-dpf wild-type and *pday* mutant retinas with zpr1 (red) and anti-rhodopsin (green) antibodies. (B’) Higher-magnification of dorsal, central, and ventral ONL of *pday* mutant retinas, which are indicated by white squares in (B). Both rhodopsin (yellow arrowheads) and zpr1 signals are observed in the peripheral region of the dorsal and ventral ONL. In addition, zpr1-positive flattened cells are located in the central ONL (white arrow). (C) TUNEL (green) of 5-dpf wild-type and *pday* mutant retinas combined with zpr1 antibody labeling (red). (C’) Higher-magnification of dorsal, central, and ventral ONL of *pday* mutant retinas, indicated by white squares in (C). Many TUNEL signals are associated with zpr1 signals (white arrows); however, a few are not associated with zpr1 (white arrowheads). (D) TUNEL (red) of 5-dpf wild-type and *pday* mutant retinas combined with anti-rhodopsin antibody labeling (green). (D’) Higher-magnification of dorsal, central, and ventral ONL of *pday* mutant retinas, indicated by white squares in (D). Rhodopsin signals are detected in the peripheral region of dorsal and ventral ONL (yellow arrowheads). TUNEL signals associated with rhodopsin signals are observed only in the peripheral region of the dorsal and ventral ONL (white arrowheads). TUNEL signals are not associated with rhodopsin signals in central ONL and non-peripheral regions of the dorsal and ventral ONL (white arrows). (E, F) Wild-type and *pday* mutant retinas with and without the transgenic line *Tg[hsp: mCherry-Bcl2]*, labelled with zpr1 (E) and zpr3 (F) antibodies. Nuclei were counterstained with Hoechst 33342 (blue). Both zpr1 and zpr3 signals in the ONL are recovered in *pday* mutant retinas with *Tg[hsp: mCherry-Bcl2]* (white arrowheads). (F’) Higher-magnification of the dorsal ONL indicated by white squares, shown in (F). Scale bars: 50 µm (A-D), 40 µm (E, F)

Next, we examined whether cones undergo apoptosis in *pday* mutants. First, we carried out double labeling of 5-dpf, wild-type, and *pday* mutant retinas with anti-zebrafish rhodopsin (Vihtelic et al., 1999) and zpr1 (Larison and Bremiller, 1990) antibodies, which label rod OSs and double-cone-type photoreceptors (red/green cones), respectively (Fig. 3B). Consistent with data of *Tg[rho:NLS-eGFP; gnat2:NLS-tdTomato]* (Fig. 2), rods were only detected in the peripheral region of the dorsal and ventral ONL of *pday* mutants (Fig. 3B’). On the other hand, cones were observed in the peripheral region of the dorsal and ventral ONL and as very flattened cells in the central ONL (Fig. 3B’). Next, we carried out TUNEL of 5-dpf, wild-type and *pday* mutant retinas combined with zpr1 antibody labeling (Fig. 3C). Many TUNEL-positive cells were observed in dorsal, central and ventral ONL of *pday* mutants (Fig. 3C’). A significant fraction of these TUNEL-positive cells was associated with zpr1 signals (Fig. 3C’). Third, we carried out TUNEL in 5-dpf, wild-type and *pday* mutant retinas combined with anti-rhodopsin antibody labeling (Fig. 3D). Many TUNEL-positive cells were observed in dorsal, central, and ventral ONL of *pday* mutants (Fig. 3D’). However, rhodopsin signals were detected only in the CMZ of the dorsal and ventral ONL (Fig. 3D’), but not in the central ONL. Thus, many TUNEL signals were not associated with rhodopsin signals in the central ONL (Fig. 3D’). Since flattened photoreceptors in the central ONL of 5 dpf *pday* mutants are cones (Fig. 2), cones undergo apoptosis at or before 5 dpf in *pday* mutants.

To confirm that both rods and cones undergo apoptosis in *pday* mutants, we combined a transgenic line *Tg[hsp:mCherry-Bcl2]*, which expresses an anti-apoptotic protein Bcl2 (Nishiwaki and Masai, 2020), with *pday* mutants and evaluated cell survival by labeling with zpr1 (Larison and Bremiller, 1990) and zpr3 (Gao et al., 2022) antibodies, which visualize double cone photoreceptors and rhodopsin/green opsin, respectively. Combined with quantitative analyses (Fig. S2A, S2B), we found that Bcl2 overexpression significantly rescued survival of both cones and rods in *pday* mutants at 6 dpf (Fig. 3E, 3F; Fig. S4D, S4E) even along the DV axis (Fig. S4F), suggesting that both cones and rods undergo apoptosis in *pday* mutants.

Although rods and cones survived in *pday* mutants that overexpressed Bcl2, zpr3 signals fail to be localized in the OS, and spread in the plasma membrane throughout the cell body, including the basal synaptic region (Fig. 3F’), suggesting that green opsin and rhodopsin are mislocalized in Bcl2-overexpressing *pday* mutant photoreceptors. Furthermore, photoreceptor nuclear shape differed between Bcl2-overexpressing *pday* mutants and wild-type siblings (Fig. S4G). Next, we examined OKR in embryos produced by crosses of *pday* heterozygous fish. On average, 25% of embryos were homozygous for the *pday* mutation and did not show OKR. Bcl2 overexpression did not restore OKR of *pday* mutants (Fig. S4H). Thus, MAK activity is primarily required for visual functions of photoreceptors, disruption of which may subsequently cause Bax-mediated apoptosis.

### *mak* is responsible for *pday* mutant phenotypes

To validate whether the *mak* gene is responsible for *pday* mutant phenotypes, we generated a zebrafish transgenic line *Tg[crx:eGFP-MAK],* which expresses the N-terminal eGFP-tagged, full-length isoform of MAK under control of the *crx* promotor. A transgenic line *Tg[crx:eGFP]* was used as a control. Labeling with zpr1 antibody revealed that *Tg[crx:eGFP-MAK]* rescued cone photoreceptor degeneration in *pday* mutants at 6 dpf, whereas *Tg[crx:eGFP]* did not (Fig. 4A, 4A’, and 4B). Furthermore, we examined OKR in embryos produced by crosses of *pday* heterozygous fish. On average, 25% of embryos were homozygous for the *pday* mutation and did not show OKR. However, *Tg[crx:eGFP-MAK]* restored OKR of *pday* mutants at 6 dpf, whereas *Tg[crx:eGFP]* did not (Fig. 4C). Thus, *mak* is responsible for *pday* mutant phenotypes: photoreceptor degeneration, and visual response defects.

**Fig. 4:**
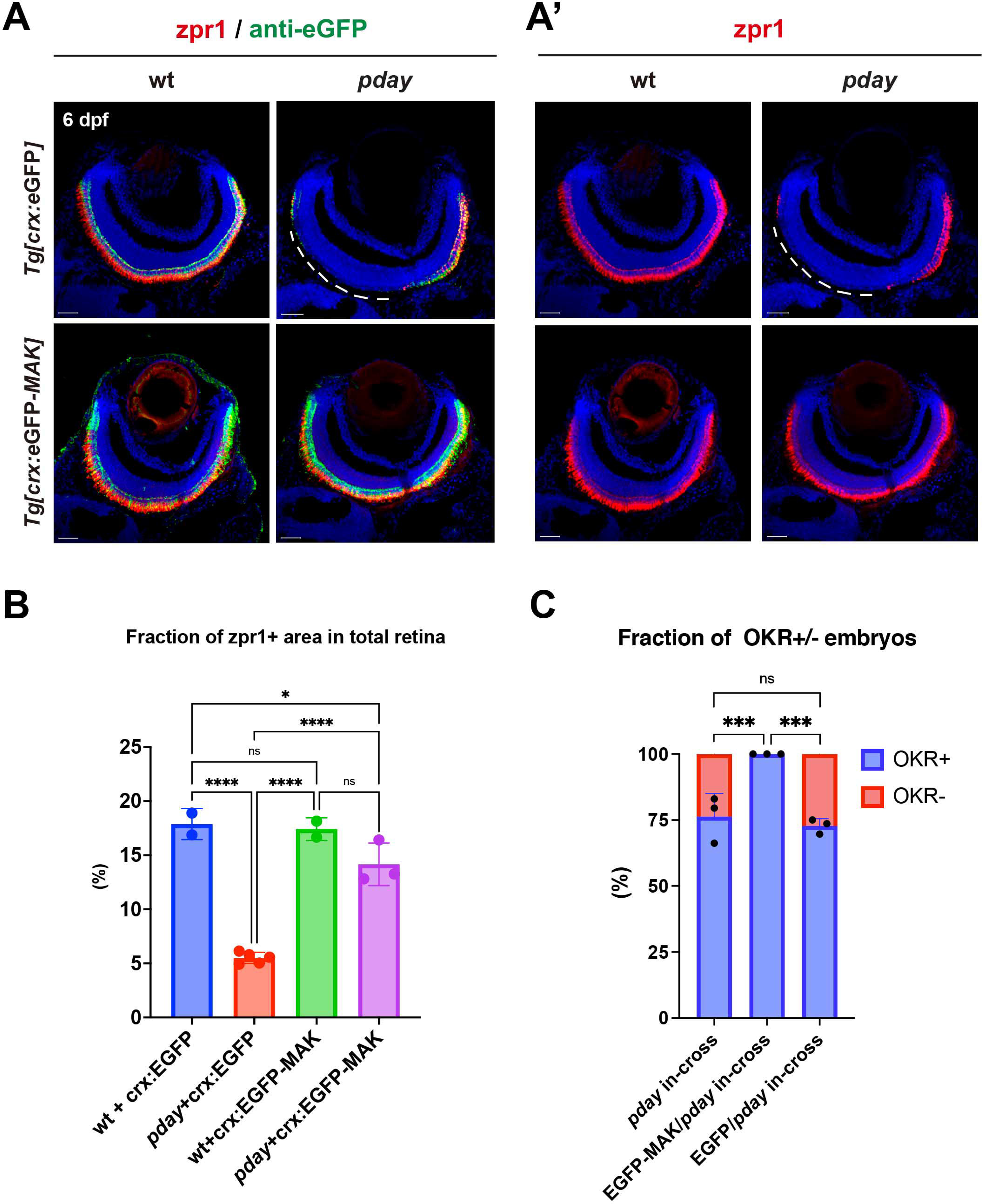
Overexpression of MAK restores photoreceptor survival and visual function of *pday* mutants. (A) Labeling of wild-type and *pday* mutant retinas with zpr1 (red) and anti-eGFP (green) antibodies carrying transgenic line *Tg[crx:eGFP]* or *Tg[crx:eGFP-MAK]*. Nuclei were counterstained with Hoechst 33342 (blue). (A’) Only red channel. zpr1 signals are markedly decreased in the dorso-central region of *pday* mutant retinas carrying *Tg[crx:eGFP]* (white dotted lines), but maintained in *pday* mutant retinas carrying *Tg[crx:eGFP-MAK]*. (B) The fraction of zpr1-positive area in the total retinal area in wild-type and *pday* mutants with *Tg[crx:eGFP]* or *Tg[crx:eGFP-MAK]*. The fraction is markedly reduced in *pday* mutants with *Tg[crx:eGFP]* (red bar), but recovered in *pday* mutants with *Tg[crx:eGFP-MAK]* (purple bar), equivalent to wild-type with *Tg[crx:eGFP]* (blue bar) or *Tg[crx:eGFP-MAK]* (green bar). Bars and lines indicate mean ± SD. Statistical significance was evaluated with ordinary one-way ANOVA and Turkey’s multiple comparison tests; *p<0.0332, ****p<0.0002, ns: not significant. (C) The fraction of OKR^+^ (blue) and OKR^-^ (red) embryos relative to total progeny produced by crosses of *pday^+/-^* parent fish, *pday^+/-^* parent fish carrying *Tg[crx:eGFP]*, and *pday^+/-^* parent fish carrying *Tg[crx:eGFP-MAK].* 57 of 244 (23.3%) embryos produced by three independent crosses of *pday^+/-^* parent fish were negative for OKR. However, 100% of total 257 embryos produced by three independent crosses of *pday^+/-^* parent fish carrying *Tg[crx:eGFP-MAK]* were positive for OKR. 68 of 266 (25.5%) embryos produced by three independent crosses of *pday^+/-^* parent fish carrying *Tg[crx:eGFP]* were negative for OKR. Bars and lines indicate mean ± SD. Statistical significance was evaluated with 2-way ANOVA and Turkey’s multiple comparison tests. *** p<0.0002, ns: not significant. Scale bars: 20 µm (A, A’)

### MAK is localized in the connecting cilium and promotes axoneme development during ciliogenesis

We examined *mak* mRNA expression during zebrafish embryonic development by whole-mount *in situ* hybridization (Fig. 5A). *mak* mRNA is ubiquitously expressed from 1- to 4-cell stages, indicating maternal mRNA expression. *mak* mRNA expression disappeared at the 50%-epiboly stage, but was detected in Kupffer’s vesicle, a ciliated embryonic organ that is important for left-right asymmetry patterning (Essner et al., 2005), at the tail-bud stage. At 24 hpf, *mak* mRNA is expressed in the central nervous system, and especially strongly expressed in the epiphysis and retina. At 48 hpf, *mak* mRNA is prominently expressed in the brain, the retina, and the pectoral fin bud. At 72 hpf, *mak* mRNA is highly expressed in the brain, including the optic tectum and the photoreceptor layer of the retina. We also confirmed that *mak* mRNA expression was markedly decreased in *pday* mutant embryos at 72 hpf, probably due to nonsense-mediated mRNA decay (Fig. S5A). Thus, *mak* mRNA is expressed in various cell types in zebrafish embryos, including Kupffer’s vesicle and retinal and pineal photoreceptors, in which ciliation supports these cell functions (Essner et al., 2005; Liu et al., 2004; Sukumaran and Perkins, 2009; Zhai et al., 2014). In addition, zebrafish mutants with cilia defects tend to develop scoliosis (Gray et al., 2021; Latour et al., 2020; Wang et al., 2022), which was observed in *pday* mutants at 2 mpf (Fig. 1A). Thus, MAK regulates cilium-associated mechanisms in photoreceptors in zebrafish.

**Fig. 5:**
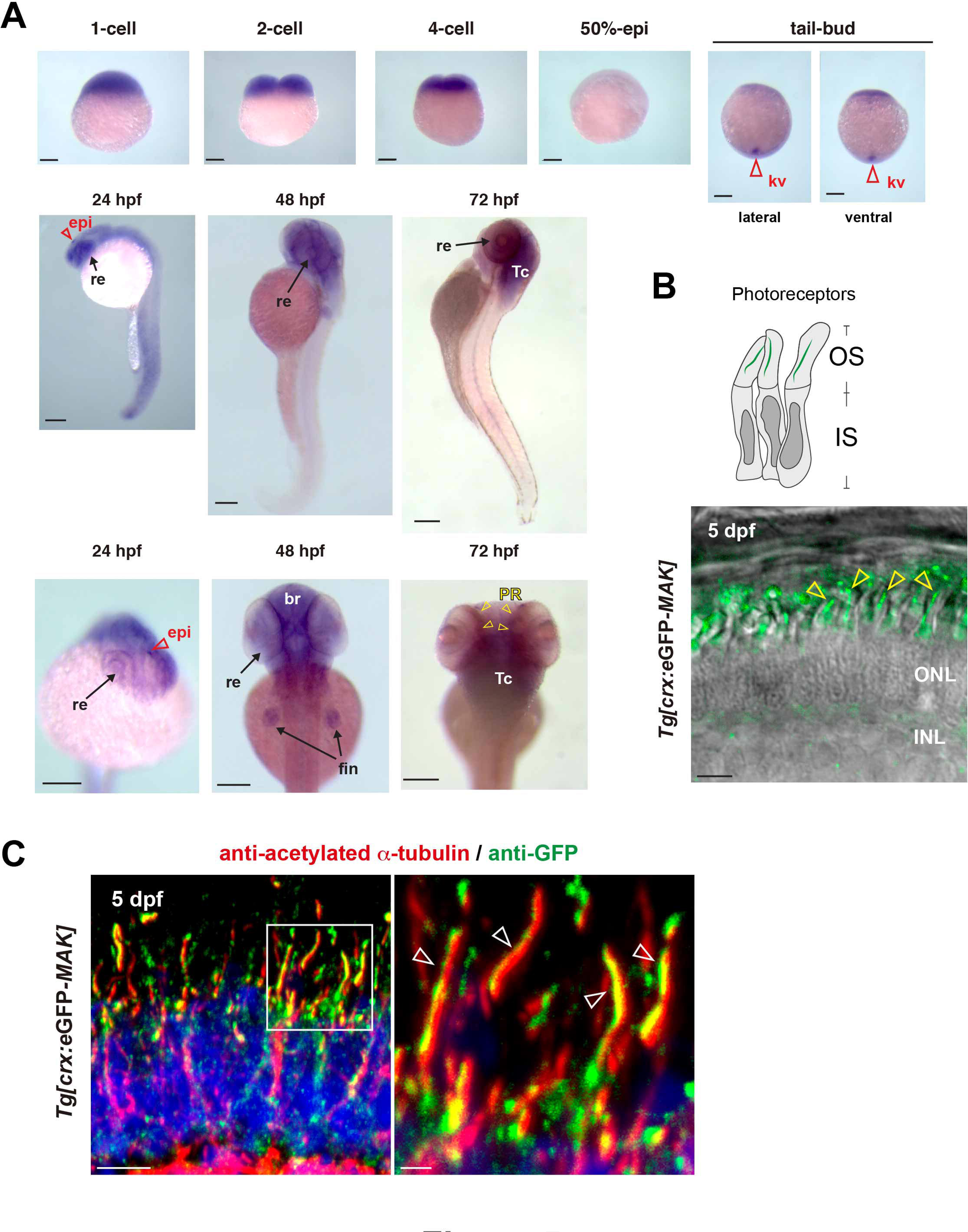
MAK is localized in axonemes of photoreceptor cilia. (A) Whole-mount *in situ* hybridization of zebrafish embryos with *mak* RNA probe. *mak* mRNA expression is observed in Kupffer’s vesicles (red open arrowhead) at the tail-bud stage, in retina (arrows) and epiphysis (red open arrowhead) at 24 hpf, and in retina, brain and fin (arrows) at 48 hpf. Retinal expression is restricted to the photoreceptor layer (yellow arrowheads) at 72 hpf. kv, Kuffper’s vesicle; re, retina; epi, epiphysis; br, brain; fin, pectoral fin; Tc, optic tectum; PR, photoreceptor layer. (B) Confocal live image of *Tg[crx: eGFP-MAK]* transgenic retina at 5 dpf. eGFP-MAK (green) is localized in the OS of photoreceptors (yellow arrowheads). (C) Labeling of 5-dpf, *Tg[crx: eGFP-MAK]* transgenic retina with anti-acetylated α-tubulin (red) and anti-GFP (green) antibodies. Nuclei were counterstained with Hoechst 33342 (blue). The right panel indicates higher magnification of the square shown in the left panel. eGFP-MAK is localized in the axoneme (white open arrowheads, right panel). Scale bars: 200 µm (A), 5 µm (B, C left), and 1 µm (C, right).

Next, we examined MAK protein localization in zebrafish photoreceptors by tracking the eGFP signal in *Tg[crx:eGFP-MAK]* retinas. eGFP-MAK is localized in a stem-like structure inside the OS of live photoreceptors (Fig. 5B). Next, we examined double labeling of *Tg[crx:eGFP-MAK]* retinas with anti-GFP antibody and anti-acetylated α-tubulin antibody. The latter mainly labels the ciliary axoneme. eGFP-MAK was co-localized to the ciliary axoneme visualized by an anti-acetylated α-tubulin antibody (Fig. 5C, Fig. S5B). Although overexpression of eGFP-MAK could lead to its localization to places where endogenous protein is not, it is likely that MAK is a ciliary protein in photoreceptors.

In zebrafish, photoreceptor precursors undergo one round of symmetric cell division to produce two daughter cones after 60 hpf, in which apically localized centrosomes start ciliogenesis (Zolessi et al., 2021). So, we examined the ciliary structure in *pday* mutant photoreceptors at 3 dpf. We used anti-γ tubulin and anti-acetylated α tubulin antibodies, which visualize the basal body and the axoneme of the cilium, respectively (Chen et al., 2021). Eyes shut (EYS) is a secreted extracellular matrix protein localized near at the transition zone of photoreceptors and required for the ciliary pocket formation in zebrafish (Yu et al., 2016). CEP290 (also known as NPHP6) is a large multidomain coiled coil protein localized at the proximal part of the transition zone (Goncalves and Pelletier, 2017; Park and Leroux, 2022). Genetic mutations of CEP290 are associated with Leber’s congenital amaurosis in humans (den Hollander et al., 2006) and photoreceptor degeneration in zebrafish (Cardenas-Rodriguez et al., 2021). So, we used anti-EYS and anti-CEP290 antibodies, which visualize transition zone-related areas. Labeling with anti-acetylated α tubulin and anti-EYS antibodies revealed that EYS signals were normally detected, but acetylated α tubulin signals were absent in *pday* mutant photoreceptors (Fig. 6A). Labeling with anti-γ tubulin and anti-EYS antibodies revealed that both γ tubulin and EYS signals were normally associated in *pday* mutant photoreceptors (Fig. 6B). Next, labeling with anti-γ tubulin and anti-CEP290 antibodies revealed that γ tubulin signals were normal in *pday* mutant photoreceptors; however, CEP290 signals were drastically reduced (Fig. 6C). These data indicate that the ciliary axoneme is absent and the transition zone is also defective in *pday* mutants, whereas the basal body seems to be normal, suggesting that the ciliary axoneme and the transition zone are the primary target of MAK. On the other hand, normal EYS signals in *pday* mutant photoreceptors suggest that MAK is not involved in EYS-mediated ciliary pocket formation.

**Fig. 6:**
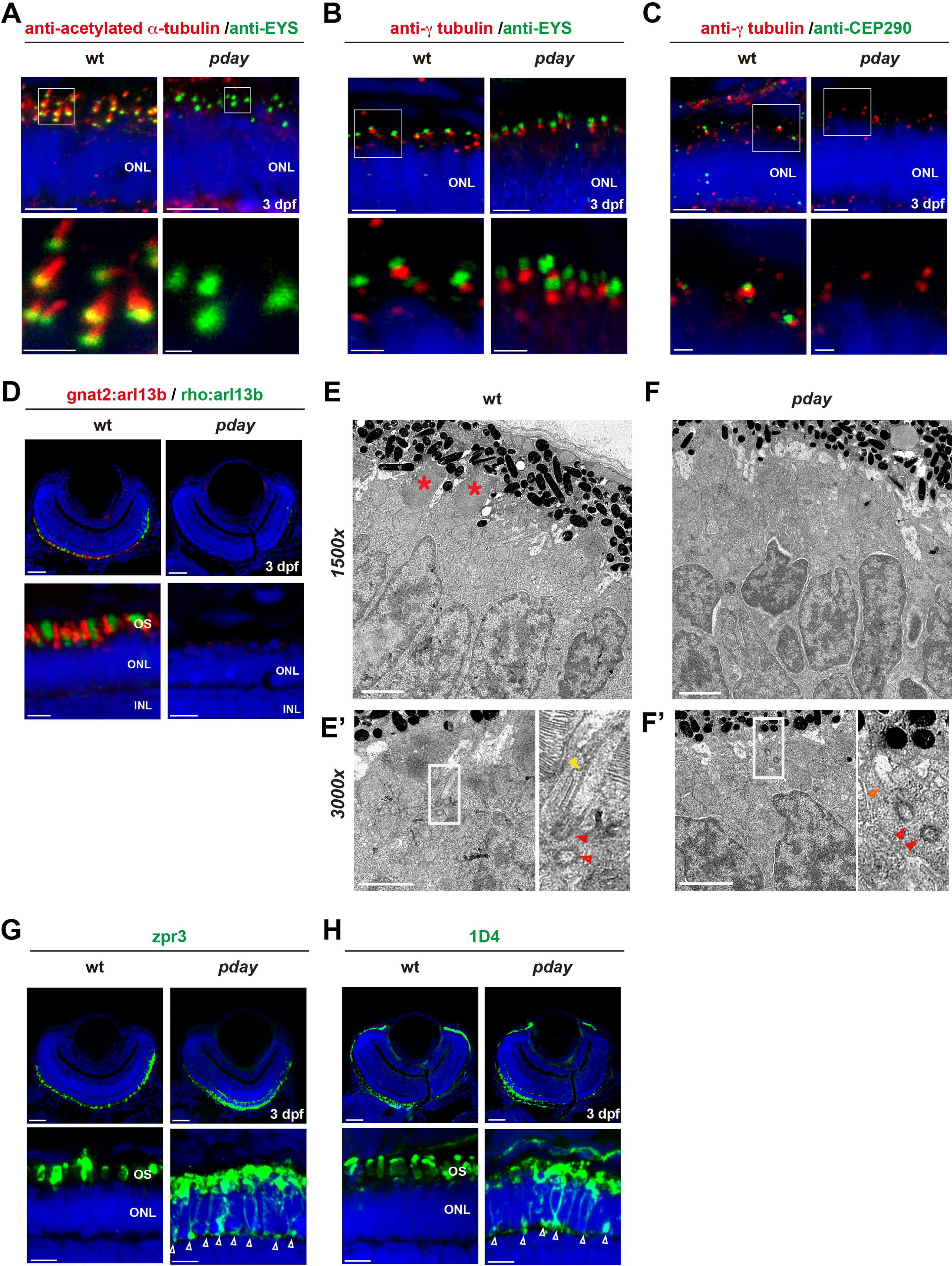
The axoneme is absent in *pday* mutant photoreceptors. (A) Wild-type and *pday* mutant retinas with anti-acetylated α-tubulin (red) and anti-EYS (green) antibodies. Nuclei were counterstained with Hoechst 33342 (blue). Bottom panels are higher magnification of the ONL shown in top panels. (B) Wild-type and *pday* mutant retinas with anti-γ-tubulin (red) and anti-EYS (green) antibodies. Nuclei were counterstained with Hoechst 33342 (blue). Bottom panels are higher magnification of the ONL shown in top panels. (C) Wild-type and *pday* mutant retinas with anti-γ-tubulin (red) and anti-CEP290 (green) antibodies. Nuclei were counterstained with Hoechst 33342 (blue). Bottom panels show higher magnification views of the ONL seen in the top panels. (D) Wild-type and *pday* mutant retinas combined with the transgenic lines, *Tg[gnat2: arl13b-tdTomato]* (red) and *Tg[rho: ail13b-eGFP]* (green). Nuclei were counterstained with Hoechst 33342 (blue). Bottom panels are higher magnification of the ONL. (E, F) EM analysis of wild-type (E) and *pday* mutant (F) photoreceptors at 3 dpf. Bottom left panels indicate higher-magnification (3000x) of apical region of photoreceptors in wild-type siblings (E’) and *pday* mutants (F’). Bottom right panels are enlarged images of the square region shown in the bottom left panels. Red asterisks indicate the OS. Red and yellow arrowheads indicate basal body and ciliary axoneme, respectively. Orange arrowheads indicate short ciliary extension from the basal body observed in *pday* mutants. (G, H) Labeling of wild-type and *pday* mutant retinas with zpr3 antibody (G) and 1D4 antibody (H). Nuclei were counterstained with Hoechst 33342 (blue). Bottom panels are higher magnification of the ONL. Arrowheads indicate ectopic distribution of zpr3 and 1D4 signals in plasma membrane including synaptic area in *pday* mutants. Scale bars: 5 µm (top panels of A, B, C; bottom panels of D, G, H), 1 µm (bottom panels of A, B, C; E, E’, F, F’), and 20 µm (top panels of D, G, H)

The OS is a specialized cilium of photoreceptors (Bachmann-Gagescu and Neuhauss, 2019; Chen et al., 2021). Arl13b is a Joubert syndrome protein, which is localized in the primary cilium and mutated in zebrafish *scorpion* (*sco*) mutants (Duldulao et al., 2009). In addition, eGFP-tagged Arl13b visualizes cilia in zebrafish (Bachmann-Gagescu et al., 2011; Borovina et al., 2010). We established two zebrafish transgenic lines, *Tg[gnat2:arl13b-tdTomato; rho:arl13b-eGFP]*, and found that fluorescent protein-tagged Arl13b overexpressed under the *gnat2* and *rho* promoters was localized more broadly within the OSs of cones and rods in wild-type retinas (Fig. 6D). However, both Arl13b-tdTomato and Arl13b-eGFP expression were absent in 3-dpf *pday* mutant photoreceptors. These data suggest that OS formation is compromised in both rod and cone photoreceptors in *pday* mutants.

Next, we applied electron microscopy (EM) to examine the ultrastructure of photoreceptors at 3 dpf (Fig. 6E, 6F). In wild-type photoreceptors at 3 dpf, photoreceptors show columnar shapes and their OSs started to form (Fig. 6E). On the other hand, OSs were not observed in *pday* mutant photoreceptors (Fig. 6F). Higher magnification EM images detected a basal body (a pair of centrioles) near the apical domain underneath the OS, as well as a long extension from the basal body, which corresponds to the transition zone and the axoneme, in wild-type photoreceptors (Fig. 6E’). However, a basal body and only a short extension from the basal body, which might correspond to part of the transition zone, were observed in *pday* mutant photoreceptors (Fig. 6F’). These data suggest that the OS and the ciliary axoneme are absent in *pday* mutant photoreceptors, whereas the basal body is normally formed. Thus, MAK regulates photoreceptor ciliogenesis by promoting axonemal formation.

### Phototransduction molecules fail to be transported to the OS in *pday* mutant photoreceptors

Phototransduction molecules are normally transported to the OS through the cilium (Athanasiou et al., 2018; Gulati and Palczewski, 2023; Karan et al., 2008; Wang and Deretic, 2014). However, mislocalization of opsins leads to photoreceptor degeneration, although detailed mechanisms are not fully understood (Alfinito and Townes-Anderson, 2002; Lopes et al., 2010; Rohrer et al., 2005). Therefore, we examined distributions of opsins and rhodopsin in *pday* mutant photoreceptors using zpr3 (Gao et al., 2022) and 1D4 (Yin et al., 2012) antibodies, which label green opsin/rhodopsin and red opsin in zebrafish, respectively. In 3 dpf wild-type photoreceptors, rhodopsin and red/green opsins were normally localized in the OS (Fig. 6G, 6H). However, in *pday* mutants, these opsins were mislocalized to the IS, plasma membrane, and the basal synaptic region (Fig. 6G, 6H). Thus, transport of phototransduction molecules to the OS is compromised in *mak* mutants.

### Kinase activity of MAK is essential for photoreceptor survival and ciliary axoneme formation

To understand how MAK regulates photoreceptor survival and ciliary axoneme formation, we focused on kinase-associated motifs (Wang and Kung, 2012) of MAK and conducted site-directed mutagenesis to silence kinase activity (Fig. 7A). Like human and mouse MAK, the kinase domain is located at the N-terminal region of zebrafish MAK (Fig. S1). In this kinase domain, KR (32-33 aa) and TDY (157-159 aa) are required for kinase activity as an ATP-binding motif and an autophosphorylation site, respectively (Wang and Kung, 2012). We designed two kinase-dead mutants of zebrafish MAK, namely AR MAK and ADF MAK, by replacing KR and TDY in wild-type zebrafish MAK with AR and ADF, respectively (Fig. 7A). We also prepared DM MAK, which carries both AR and ADF mutant motifs. *In vitro* kinase assay using MAK proteins purified from an *E.* coli. expression system revealed that all three kinase-dead MAK mutants, AR MAK, ADF MAK, and DM MAK, failed to phosphorylate a MAK mock substrate, myelin basic protein (MBP), indicating that MAK kinase activity is silenced in either mutant, AR MAK, ADF MAK, or DM MAK (Fig. 7B).

**Fig. 7:**
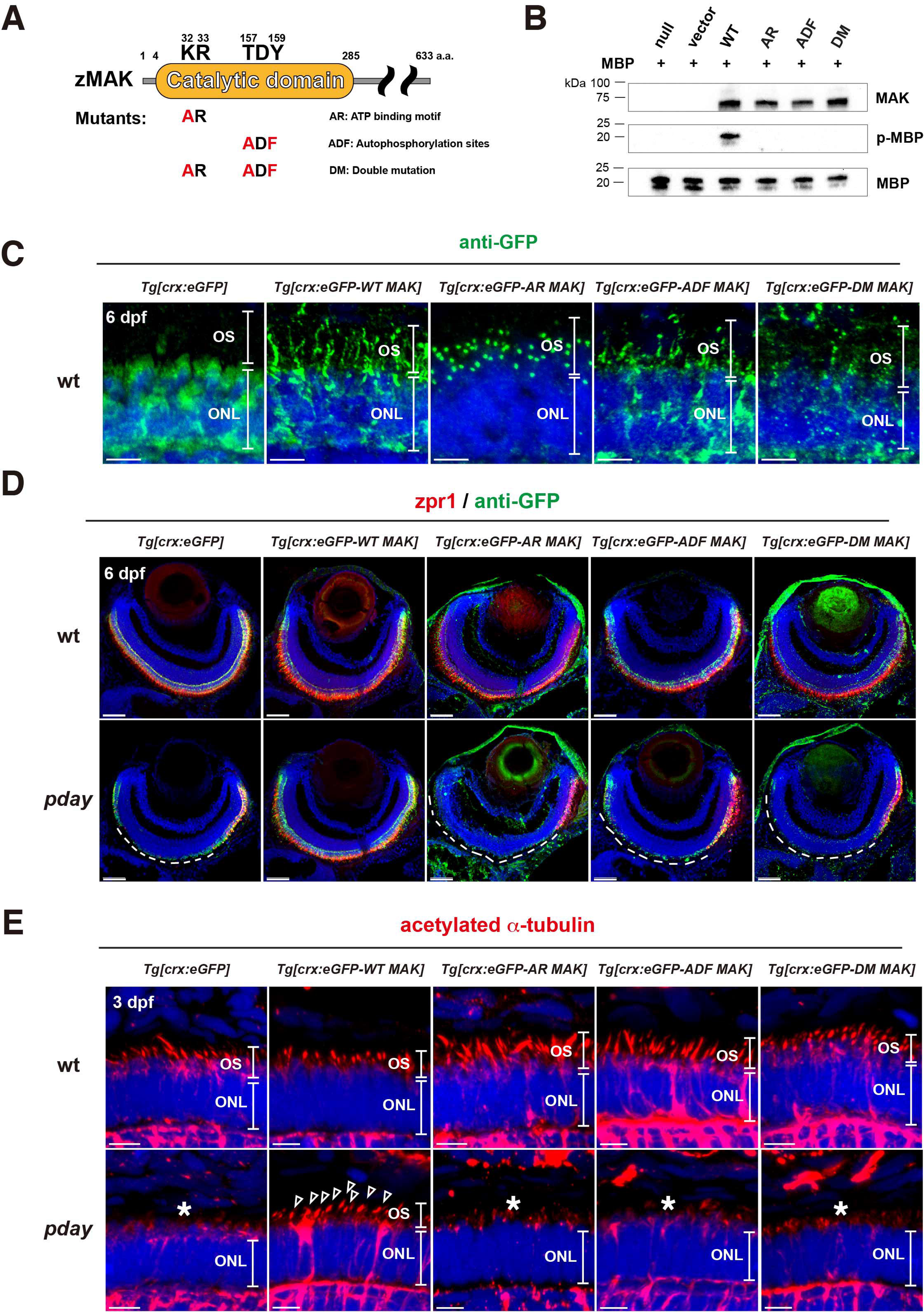
MAK kinase activity is required for MAK protein localization in the ciliary axoneme, axoneme formation, and photoreceptor maintenance. (A) A scheme of generation of three types of MAK kinase-dead mutants: AR MAK, ADF MAK and DM MAK. Zebrafish MAK has a catalytic domain, which contains an ATP binding motif, KR, and autophosphorylation site, TDY. AR MAK and ADF MAK were generated by conversion of KR and TDY into AR and ADF, respectively. DM MAK has both AR and ADF mutations. (B) *In vitro* kinase assay of MAK using a substrate, MBP. Only wild-type MAK phosphorylates MBP. (C) Labeling of 6-dpf, wild-type ONL carrying the transgenic line, *Tg[crx:eGFP], Tg[crx:eGFP-WT MAK], Tg[crx:eGFP-AR MAK], Tg[crx:eGFP-ADF MAK]*, or *Tg[crx:eGFP-DM MAK]*, with anti-GFP antibody (green). Nuclei were counterstained with Hoechst 33342 (blue). (D) Labeling of 6-dpf, wild-type and *pday* mutant retinas carrying a transgenic line, *Tg[crx:eGFP], Tg[crx:eGFP-WT MAK], Tg[crx:eGFP-AR MAK], Tg[crx:eGFP-ADF MAK],* or *Tg[crx:eGFP-DM MAK]*, with zpr1 (red) and anti-GFP (green) antibodies. Nuclei were counterstained with Hoechst 33342 (blue). zpr1-positive photoreceptors are degenerated in dorso-central retinas of *pday* mutants carrying *Tg[crx:eGFP]* (white dotted line). However, photoreceptor degeneration is rescued only in *pday* mutants carrying *Tg[crx:eGFP-WT MAK]*, but not in *pday* mutants carrying kinase-dead mutant transgenes (white dotted line). (E) Labeling of 3-dpf, wild-type and *pday* mutant ONL carrying the transgenic line, *Tg[crx:eGFP], Tg[crx:eGFP-WT MAK], Tg[crx:eGFP-AR MAK], Tg[crx:eGFP-ADF MAK],* or *Tg[crx:eGFP-DM MAK]*, with anti-acetylated α-tubulin antibody (red). Nuclei were counterstained with Hoechst 33342 (blue). Axoneme formation defects are rescued only in *pday* mutants carrying *Tg[crx:eGFP-WT MAK]* (arrowheads), but not in *pday* mutants carrying kinase-dead mutant transgenes (asterisks). Scale bars: 5 µm (C, E) and 40 µm (D).

Next, we generated zebrafish transgenic lines, *Tg[crx:eGFP-AR MAK]*, *Tg[crx:eGFP-ADF MAK]*, and *Tg[crx:eGFP-DM MAK]*, by introducing kinase-dead mutant motifs into the wild-type MAK-expressing construct, *Tg[crx:eGFP-WT MAK]*. First, we examined subcellular localization of these three kinase-dead mutant MAK proteins in zebrafish photoreceptors by labeling these transgenic wild-type retinas with anti-GFP antibody (Fig. 7C). eGFP-WT MAK was localized in a stem-like pattern similar to the axoneme of photoreceptor cilium. However, eGFP-AR MAK was localized in a spot-like region near the proximal region of photoreceptor cilium, which seems to correspond to either the basal body or the transition zone. Thus, ATP-binding activity is important for proper localization of MAK in the axoneme of photoreceptor cilium. eGFP-ADF MAK was likely to be localized in the axoneme of photoreceptor cilium, but its expression level was low, suggesting that eGFP-ADF MAK may be unstable. eGFP-DM MAK showed low expression level as well, which made it difficult to define its subcellular localization. Thus, MAK kinase activity influences its localization in the axoneme of photoreceptor cilium.

Next, using zpr1 antibody, we examined whether MAK kinase activity is required for survival of double-cone photoreceptors. We combined transgenic lines, *Tg[crx:eGFP-AR MAK]*, *Tg[crx:eGFP-ADF MAK]*, and *Tg[crx:eGFP-DM MAK]*, with the *pday* mutants. In contrast to *Tg[crx:eGFP-WT MAK]*, all three kinase-dead mutant MAK transgenes failed to rescue survival of double-cone photoreceptors at 6 dpf (Fig. 7D), suggesting that MAK kinase activity is important for photoreceptor survival. Furthermore, we examined ciliary axonemes in *pday* mutants by labeling *pday* mutants combined with *Tg[crx:eGFP-AR MAK]*, *Tg[crx:eGFP-ADF MAK]*, and *Tg[crx:eGFP-DM MAK]* with anti-acetylated α-tubulin antibody. In contrast to *Tg[crx:eGFP-WT MAK]*, all three kinase-dead mutant MAK transgenes failed to rescue axoneme formation at 6 dpf (Fig. 7E). Thus, MAK kinase activity is critical for ciliary axoneme development and photoreceptor survival (Fig. 8).

**Fig. 8:**
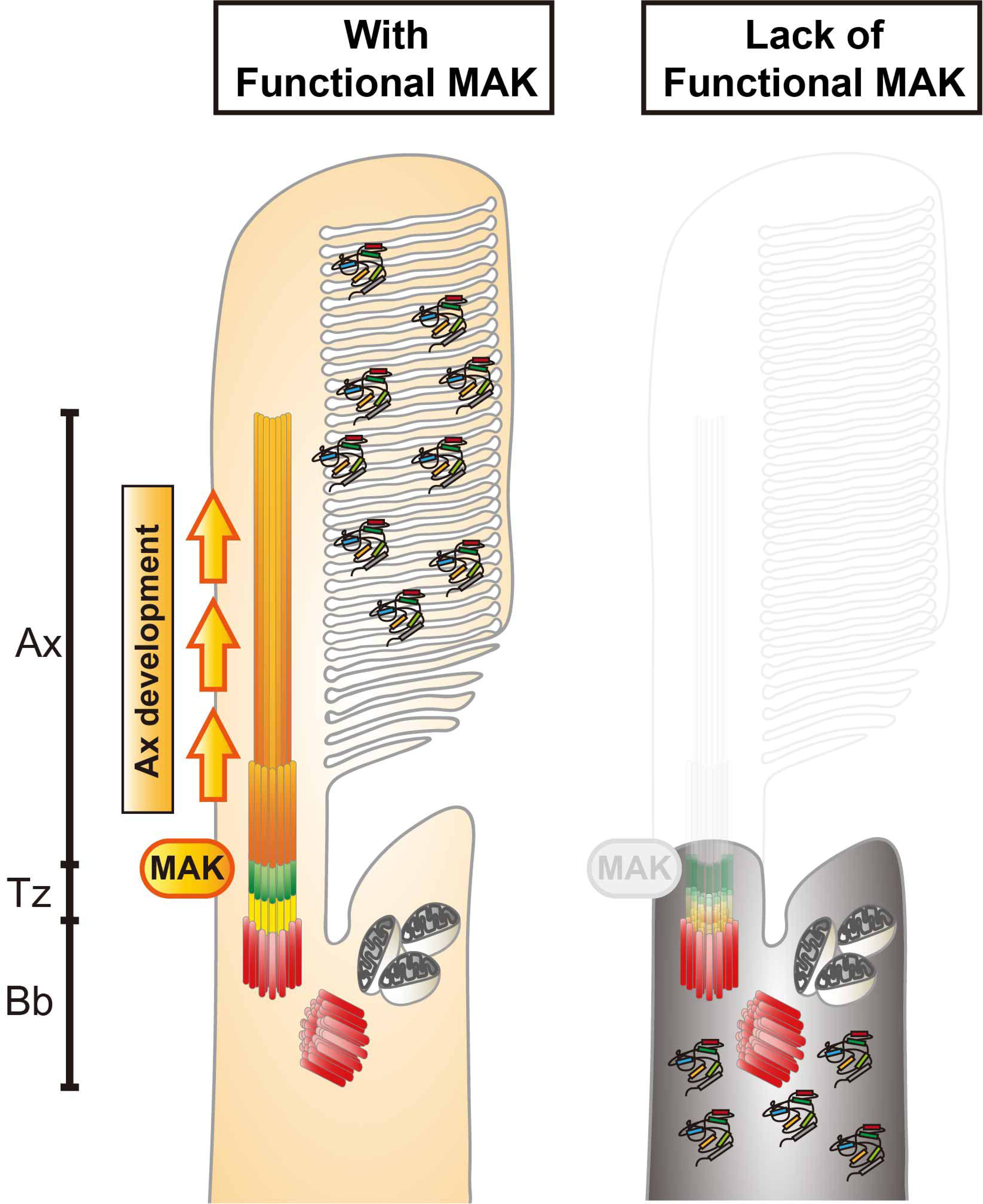
MAK is essential for ciliary axoneme formation in zebrafish photoreceptor ciliogenesis. Phenotypes of zebrafish *mak* mutant photoreceptors. In the absence of MAK, the axoneme (orange) does not form and the transition zone (green) is affected, resulting in failure to form the OS and to transport phototransduction molecules into the OS.

## Discussion

In humans and other animal models including zebrafish, many genetic mutations showing defects in photoreceptor ciliogenesis have been identified (Chen et al., 2021). In zebrafish mutants of transition zone components, selective transport of phototransduction molecules into the OS, is compromised, resulting in ectopic distribution of these OS-resident proteins outside the OS (Wang et al., 2022). However, the axoneme itself is formed in almost all of these transition-zone-defective mutants. In zebrafish mutants of BBS components, which cooperate with IFT to regulate intra-ciliary transport (Tian et al., 2023), OS non-resident proteins are mislocalized into the OS, but the axoneme and the OS are formed (Masek et al., 2022). Furthermore, in zebrafish *arl13b* mutants, with compromised cargo release in the OS, the axoneme and the OS are reduced in size, but still formed (Song et al., 2016). Thus, most ciliogenesis-defective mutants show formation of axoneme structure in photoreceptors. However, there are a few exceptions. In zebrafish mutants of an IFT-B component, IFT88 (also named *oval*), Traf3ip1 (also named *elipsa*), and a Kinesin II family protein Kif3a, the axoneme and the OS are totally absent in photoreceptors (Krock and Perkins, 2008; Omori et al., 2008; Sukumaran and Perkins, 2009; Zhu et al., 2021). Interestingly, in *ift88* and *traf3ip1* mutants, a single spot-like γ-tubulin or centrin signal is detected in the apical region of photoreceptors, similar to zebrafish *mak* mutants, suggesting that basal bodies are formed in both mutants (Omori et al., 2008; Sukumaran and Perkins, 2009; Tsujikawa and Malicki, 2004). Thus, it is likely that MAK cooperates with anterograde IFT complexes including IFT88 to promote axoneme formation during ciliogenesis.

Overexpression of wild-type MAK recovers formation of axonemes, leading to photoreceptor survival, in *mak* mutants. However, overexpression of kinase-mutant forms of MAK fail to rescue defects in axoneme and cell survival, so phosphorylation events on MAK kinase substrates are necessary for axoneme formation. Interestingly, GFP-tagged AR MAK is localized in a single spot area near the apical region of photoreceptors, which seems to correspond to the basal body or transition zone, whereas GFP-tagged, wild-type MAK is located along the axoneme. One possibility is that target proteins of MAK-mediated phosphorylation exist near the basal body or transition zone and move with MAK along the axoneme after phosphorylation by MAK. Another possibility is that MAK interacts with adaptor proteins that are localized in the basal body or transition zone to support MAK substrate phosphorylation for ciliogenesis. At present, MAK phosphorylation substrates and adaptor proteins that facilitate MAK substrate phosphorylation are unknown in zebrafish; however, since the IFT complex is assembled and transported through the transition zone in the first step of ciliogenesis (Breslow and Holland, 2019), IFT complex components may be promising candidates to promote MAK substrate phosphorylation.

MAK proteins share a highly conserved amino acid sequence between zebrafish and mice. However, in contrast to zebrafish *mak* mutants, the axoneme is elongated in mouse *Mak* mutants (Omori et al., 2010). Why does *mak* gene knockout induce such opposite phenotypes in zebrafish and mice? From a phylogenetic point of view, MAK belongs to the MAK/ICK/MOK serine/threonine kinase family, all members of which share a highly conserved kinase domain (Chen et al., 2013; Miyata and Nishida, 1999). CILK1 (previously known as ICK) is a member of this kinase family and is now classified as a paralog of MAK. Indeed, human CILK1 shows a highly similar amino acid sequence in the kinase domain to that of zebrafish MAK (81.5% amino acid identity). Interestingly, in *Cilk1* mouse mutants, the ciliary axonemes of fibroblasts are shortened compared with those of wild-type controls (Chaya et al., 2014). Furthermore, mouse CILK1 interacts with IFT-B components, including IFT88 (Nakamura et al., 2020). Since a CILK1 homologous gene has not been identified in the zebrafish genomic database, zebrafish MAK and mouse CILK1 may share a common regulatory pathway for axoneme formation during ciliogenesis. It will be important to investigate whether zebrafish MAK cooperates with IFT-B to regulate axoneme formation during ciliogenesis.

Another interesting finding is that there is a difference in degeneration of rods and cones in *mak* mutants. In *mak* mutants, rods undergo acute apoptosis from 3 to 4 dpf, whereas cones undergo progressive shrinkage from 4 to 6 dpf. Interestingly, early pyknotic nuclei and later progressive shrinkage of the photoreceptor layer was also observed in zebrafish *ift88* and *traf3ip1* mutants, both of which show defects in axoneme formation (Bahadori et al., 2003; Doerre and Malicki, 2002; Sukumaran and Perkins, 2009; Wang et al., 2022). Recently, it was reported that photoreceptor degeneration in ciliogenesis-defective zebrafish *Tulp1* mutants is associated with pathological features of ferroptosis (Jia et al., 2022), which is a new form of cell death linked to accumulation of lipid peroxides (Yu et al., 2017). Although both rod and cone degeneration in *mak* mutants depend on apoptosis, it is interesting to examine why cones show progressive shrinkage in the absence of axoneme formation. Recent metabolic transcriptome analysis revealed that cellular metabolism of rods and cones depends on aerobic glycolysis and oxidative phosphorylation, respectively, but that cones shift toward glycolysis in the early pathological stage of retinitis pigmentosa, suggesting an intrinsic difference of metabolic regulation between rods and cones (Lee et al., 2023). It is interesting to examine how metabolic states regulated differently in rods and cones in *mak* mutants.

Lastly, *mak* mutants show scoliosis and smaller body size in later development. A comprehensive analysis of zebrafish transition zone mutants revealed phenotypic variations between these ciliary-defective mutants (Wang et al., 2022). Among them, *cep290* mutants show severe scoliosis in the adult stage, whereas *tmem216* mutants show smaller body size. Indeed, CEP290 is markedly decreased in *mak* mutants, so it is possible that scoliosis and smaller body size in *mak* mutants are caused by reduced activity of CEP290 and Tmem216, respectively. Further study on the ciliary regulator network in *mak* mutants will reveal the role of MAK in ciliopathy mechanism.

## Materials and Methods

### Ethics statement

Zebrafish were maintained at 28.5°C, on a 14h light/10h dark cycle, following standard procedures (Westerfield, 2000). Collected embryos were cultured in E3 embryo medium (5 mM NaCl, 0.17 mM KCl, 0.33 mM CaCl_2_, 0.33 mM MgSO_4_) containing 0.003% 1-phenyl-2-thiouera (PTU) to prevent pigmentation and 0.0002 % methylene blue to prevent fungal growth. All experiments were performed on zebrafish embryos between the one-cell stage and 6 dpf prior to sexual differentiation. Therefore, sexes of embryos could not be determined. All zebrafish experiments were performed under the OIST Animal Care and Use Program based on the Guide for the Care and Use of Laboratory Animals by the National Research Council of the Nation Academies and approved by the Association for Assessment and Accreditation of Laboratory Animal Care (AAALAC). The OIST Institutional Animal Care and Use Committee approved all experimental protocols (Protocol: ACUP-2023-016, ACUP -2023-017, ACUP -2023-018, ACUP -2023-019, ACUP -2023-020).

### Zebrafish strains and maintenance

Okinawa wild-type (oki) was used as a wild-type strain to maintain all mutant and transgenic strains. The zebrafish mutant, *pday*^s351^, was originally isolated by the Herwig Baier lab (Muto et al., 2005). Mapping of the *pday*^s351^ mutation was carried out in the genetic background of WIK. Zebrafish transgenic lines, *Tg[gnat2:NLS-tdTomato]*^oki070^ and *Tg[rho:NLS-eGFP]*^oki071^, were used to visualize cone and rod nuclei, respectively. *Tg[hsp:mCherry-Bcl2]^oki029^* (Nishiwaki and Masai, 2020) was used to express mCherry-tagged zebrafish Bcl2 under control of the zebrafish *heat-shock 70* (*hsp70*) promoter. *Tg[crx:eGFP]*^oki072^, *Tg[crx:eGFP-WT MAK]*^oki073^, *Tg[crx:eGFP-AR MAK]*^oki074^ and *Tg[crx:eGFP-ADF MAK]*^oki075^, and *Tg[crx:eGFP-DM MAK]*^oki076^ were used to express eGFP-tagged wild-type and kinase-deficient mutants MAK, respectively. *Tg[gnat2:arl13b-tdTomato]*^oki077^ and *Tg[rho:arl13b-eGFP]*^oki078^ were used to visualize cone and rod OSs, respectively.

### Mapping and cloning of the *pday* mutant gene

*pday*^+/-^ fish maintained in the *oki* wild-type background were outcrossed with the *WIK* wild-type strain to generate F1 *pday*^+/-^ fish with an *oki* and *WIK* trans-heterozygous wild-type background. These F1 *pday*^+/-^ male and female fish were crossed to produce F2 embryos. F2 *pday*^−/−^ embryos were selected by measuring OKR at 5 dpf and stored in methanol at -80°C. Genomic DNA was extracted from individual F2 *pday*^−/−^ embryos. First, two DNA pools of 20 wild-type and 20 homozygous mutant embryos, respectively, were used for PCR-mediated amplification of SSLP markers (Knapik et al., 1998; Shimoda et al., 1999) to examine which chromosome hosts the *pday* mutation. Furthermore, 93 F2 *pday*^-/-^ embryos (186 meiosis) were used to restrict the cytogenetic position of the *pday* mutation using SSLP markers mapped on the linked chromosome. We restricted the *pday* mutation within the genomic region flanked between two SSLP markers, z23011 and z13695, which were annotated at positions 8.517 and 9.123 Mb on chromosome 24 (zebrafish genomic database, GRCz10, Ensemble release 80). This region contained 10 genes (*tfap2a*, *tmem14ca*, *mak*, *gcm2*, *elovl2*, *BX546453*, *gnal*, *mppe1*, *CR318624*, and *dlgp1b*). Since, among these, *mak* is the only gene whose mutations cause inherited photoreceptor degeneration diseases, such as retinitis pigmentosa in humans (Ozgul et al., 2011; Stone et al., 2011), we designed a new SSLP marker, namely Mak-N1, which was located around 20 bp upstream of the first exon of the *mak* gene, and confirmed no recombination of Mak-N1 in all 93 F2 *pday*^-/-^ embryos (186 meiosis). Sequence information on forward and reverse primers of z23011, z13695, and Mak-N1 is provided in the appendix.

To confirm that *pday* gene encodes MAK, full-length *mak* cDNA was amplified by PCR using mRNA prepared from wild-type and *pday*^s351^ homozygous embryos at 6 dpf. However, only a partial cDNA fragment that corresponds to the C-terminal coding region covering from the middle of exons 7 to 15 was amplified by PCR from *pday* homozygous mutant mRNA. We sequenced the full-length cDNA of wild type and the C-terminal partial cDNA of *pday* homozygous mutants and found no amino acid change in the MAK coding region from exons 8 to 15 between wild-type and *pday^-/-^* cDNA. Next, we amplified six DNA fragments from *the pday^-/-^* genome, each of which covers six N-terminal coding exons (the exon 2 - 7), respectively. Sequencing of these six DNA fragments revealed a non-sense mutation in exon 4 of the *pday^-/-^* genome. To confirm that the *pday* mutant gene encodes MAK, we combined *pday* mutants with the transgenic line *Tg[crx:eGFP-WT MAK]*^oki073^, and showed that expression of eGFP-tagged MAK in photoreceptor precursors rescues photoreceptor degeneration in *pday* mutants.

### Generation of DNA expression constructs and their transgenic lines

*Tol2[crx: membrane targeted YFP (MYFP)]* was used as a template to make the expression construct *Tol2[crx: eGFP-MAK]*. *Tol2[crx: MYFP]* was constructed using the To2kit multiple gateway-based construction system (Kwan et al., 2007), whose Tol2 vector backbone contains the *cmcl2:CFP* transgenesis marker (Suzuki et al., 2013). *mak* cDNA corresponding to a full-length isoform annotated on the zebrafish genomic database (mak-201, GRCz11, Ensemble release 110), was amplified by PCR using a mRNA pool of 3-dpf, wild-type embryos and was subcloned into pCR TOPO vector (Invitrogen). After the eGFP cDNA fragment was tagged in-frame to the *mak* full-length cDNA at the N-terminus, the MYFP region of *Tol2[crx: MYFP]* was replaced with the DNA fragment encoding eGFP-tagged *mak* cDNA at the *BamHI* and *ClaI* sites to make the expression construct *pTol2[crx:eGFP-WT MAK]*. The *pTol2[crx:eGFP-WT MAK]* was used as a template in site-directed mutagenesis for making three kinase-dead MAK constructs, *Tg[crx:eGFP-AR MAK]*, *Tg[crx:eGFP-ADF MAK]* and *Tg[crx:eGFP-DM MAK]*.

The *EF1α* promoter of the Tol2 transposon vector pT2AL200R150G (Urasaki et al., 2006) was replaced with the zebrafish *gnat2* promoter (Iribarne et al., 2019) at the *XhoI* and *BamHI* site to make *Tol2[gnat2:eGFP]*. Zebrafish *arl13b* cDNA was amplified by PCR using an mRNA pool prepared from 4-dpf, wild-type embryos and subcloned into the pCR TOPO vector (Invitrogen). After a tdTomato cDNA fragment was tagged in-frame to the *arl13b* cDNA at the C-terminus, a DNA fragment of tdTomato tagged *arl13b* cDNA was further subcloned into the modified pT2AL200R150G carrying the *gnat2* promoter at the *BamHI* and *ClaI* sites, to make the expression construct *pTol2[gnat2:arl13b-tdTomato]*. Similarly, the *EF1α* promoter of the Tol2 transposon vector pT2AL200R150G was replaced with the zebrafish *rhodopsin* promoter (1.1kb upstream of the transcription unit) at the *XhoI* and *BamHI* site. A DNA fragment of C-terminal eGFP-tagged *arl13b* cDNA was prepared and subcloned into the modified pT2AL200R150G carrying the *rhodopsin* promoter at the *BamHI* and *ClaI* sites, to make the expression construct *pTol2[rho:arl13b-eGFP]*.

The nuclear localization signal (NLS) was fused to cDNA encoding tdTomato or eGFP at the N-terminus. DNA fragments of NLS-tdTomato and NLS-eGFP were subcloned into the modified pT2AL200R150G carrying the zebrafish *gnat2* promoter and the zebrafish *rhodopsin* promoter, respectively, at the *BamHI* and *ClaI* sites, to make the expression constructs *pTol2[gnat2:NLS-tdTomato]* and *pTol2[rho:NLS-eGFP]*.

Next, these Tol2 expression construct plasmids and Tol2 transposase mRNA were co-microinjected into one-cell-stage zebrafish fertilized eggs for generation of transgenic lines, *Tg[crx:eGFP-WT MAK], Tg[crx:eGFP-AR MAK], Tg[crx:eGFP-ADF MAK], Tg[crx:eGFP-DM MAK], Tg[gnat2:arl13b-tdTomato]*, *Tg[rho:arl13b-eGFP], Tg[gnat2:NLS-tdTomato]* and *Tg[rho:NLS-eGFP]*. Injected F0 fish expressing fluorescent protein (eGFP or tdTomato) in the ONL were bred until adulthood and used to identify founder fish, which produce fluorescent protein-expressing F1 progeny. These transgenic F1 embryos were raised and used to produce F2 generation for establishment of the transgenic strain.

### Histology

For plastic sections, zebrafish embryos were anesthetized in 0.02% 3-aminobenzoic acid ethyl ester (tricaine) and fixed in PBS (150mM NaCl, 10mM PO4^3-^, pH 7.4) (Invitrogen) with 4% paraformaldehyde (PFA) at 4°C overnight. Samples were dehydrated with an ethanol gradient before being embedded in JB4 resin (Polysciences). Samples were sectioned at 5 µm with a microtome Rotatif, NM335E (MICROM International GmbH) and stained with 0.1% toluidine blue for further analysis.

For TUNEL, samples were covered with the staining mixture at 37°C for 1 h following instructions of the In Situ Cell Death Detection Kit (Roche, 11684795910). After the TUNEL reaction, nuclei were stained with 60 mM PO_4_ buffer (PB) (42.6 mM NaH_2_PO_4_, 17.4 mM Na_2_HPO_4_, pH 7.3) containing Hoechst 33342 (Fujifilm, 346-07951) at 1:1000, followed by washing with PB with 0.1% TritonX-100 (0.1% PBTx) for 3 x 5 min. Samples were mounted with Fluoromount (Diagnostic BioSystems, K024) for further analysis.

For immunohistochemistry, 3-6-dpf embryos were fixed with 4% PFA at room temperature for 2 hours, and transferred to 30% sucrose in PB overnight before being embedded and frozen in OCT compound. However, we used non PFA-fixed samples for anti-GFP labeling shown in Fig. 6C, because the eGFP-MAK signals are stably detected only in non-fixed tissues. Cryosections were prepared at 7 µm with a cryostat (Cryostar NX70, Thermo Scientific.), and at 20 µm for imaging cilia components. Cryosections were air-dried for at least 2 h, then rehydrated in 0.5% PBTx for 3 x 5 min. Rehydrated sections were immersed in blocking solution (10% goat serum in 0.5% PBTx) for 1 h at room temperature. Then primary antibody in blocking solution at an appropriate dilution (see appendix) was applied, and sections were washed with 0.5% PBTx 3 x 10 min. Next, primary antibody-labeled sections were treated with fluorophore-conjugated secondary antibody at 1:500 and Hoechst 33342 at 1,000 (DOJINDO Laboratories, CAS23491-52-3) in blocking solution for 1 h at room temperature, or 50 nM Sytox-Green (Molecular Probes) in 0.1% PBTx 3 x 10 min after secondary antibody, and then washed with 0.1% PBTx 3 x 10 min. Sections were finally mounted with Fluoromount. Confocal images were scanned with a confocal laser-scanning microscope, LSM780 (Carl Zeiss) or Fluoview FV3000 (Olympus, Evident). Information on primary and secondary antibodies is provided in the appendix.

For EM analysis, 3-dpf embryos were fixed in PB with fixative (2.5% glutaraldehyde, 1% paraformaldehyde, 3% sucrose, and 30 mM HEPES) on ice for 2 h, followed by washing with PB with 3% sucrose and 30 mM HEPES 3 x 5 min. Embryos were postfixed in 60 mM PB with 1% OsO_4_ for 1 h on ice, followed by washing with Milli-Q water three times. Samples were dehydrated with a gradient series of ethanol that was replaced with acetone. After treatment with a gradient series of propylene oxide (PO), epoxy resin EPON812 (Nisshin-EM, 3402) was applied to the samples before they were embedded in resin and solidified for 2 days at 60°C. Ultra-thin sections at 50 nm were prepared with a Leica UC7 ultra-microtome and processed with Reynolds’ stain (Reynolds, 1963). Images were observed and captured using a JOEL JEM-1230R transmission EM.

### Bcl2 overexpression in *pday* mutant retinas

Transgenic line *Tg[hsp:mCherry-Bcl2]* (Nishiwaki and Masai, 2020) was combined with *pday* mutants. Heat shock treatment was carried out by incubation of embryos at 39°C for 1 h every 12 h after 1 dpf until 6 dpf.

### *In situ* hybridization

Whole-mount *in situ* hybridization was carried out using a published protocol (Thisse and Thisse, 2008). A DNA fragment containing *mak* full-length cDNA combined with the T7 RNA polymerase promoter, which is located at the 3’ end of the *mak* full-length cDNA, was amplified by PCR. Digoxigenin (DIG)-labeled anti-sense *mak* RNA probe was generated by *in vitro* transcription from this T7 promoter: *mak* cDNA fragment as the template using a DIG RNA labeling kit (Roche, 11175025910). Zebrafish embryos from the 1-cell stage to 72 hpf were fixed with 4% PFA at 4°C overnight, and were stored in 100% methanol at -70°C. Samples were rehydrated from methanol to PBS, and then treated with proteinase K (10 μg/mL) to increase permeability. After refixing with 4% PFA for 20 min, samples were soaked with prehybridization buffer at 65°C for 2 h, then incubated with hybridization buffer containing *mak* antisense RNA probe at 65°C overnight. Samples were washed with 2xSSC with 0.1 % Tween20 for 5 min, room temperature, and then 0.2xSSC with 0.1 % Tween20 3 x 30 min at 65°C. Washed samples were transferred to maleic acid buffer with 0.1% Tween20 (MABT), and then to MABT with 2% blocking reagent (Roche, 11096176001) at 4°C overnight. Samples were incubated with anti-DIG-AP antibody (Roche, 11093274910) at 1:6,000 in MABT for 1 h at room temperature, washed with MABT 3 x 15 min, and then transferred to Tris-HCl buffer (pH 9.5) for 15 min. For the alkaline phosphatase (AP) reaction, samples were transferred to BM-Purple (Roche, 11442074001) until a blue signal developed, and were then fixed with 4% PFA overnight at 4°C. These samples were mounted in 70% glycerol-PBS and stored at 4°C for long-term preservation. Images of the embryos were observed with a SteREO Discovery V12 dissection microscope (Carl Zeiss).

### Live imaging

For live imaging using confocal LSM, embryos were anesthetized in 0.02% tricaine and mounted on glass slides in appropriate positions and glued with 3% methylcellulose. An upright confocal LSM, FV3000 (Olympus, Evident) was used for scanning fluorescent signals in the photoreceptor OS of *Tg[crx:GFP-MAK]*.

### *In vitro* kinase assay

The wild-type zebrafish *mak* cDNA fragment was introduced into pGEX6p3 (GE Health Care Life Science). Kinase-dead *mak* cDNA fragments were generated from pGEX6p3-zMAK(wt) with primers for site-directed mutagenesis. Plasmids were transformed into *E*. *coli* strain BL21-competent cells (TaKaRa Bio, TKR9126). Colonies were inoculated into LB broth for starter cultures incubated at 37°C overnight, followed by amplification cultures containing 1% of the starters. Amplification cultures were cooled on ice once the O.D. reached 0.6 and IPTG was added (TaKaRa Bio, TKR9030) to a final concentration of 1 mM, and incubated at 20°C overnight. Cultures were pelleted, washed with cold PBS, and sonicated in 1% PBS TritonX-100 (PBSTx) with 1x protease inhibitor (Roche, 04693132001). Debris was then pelleted, and supernatant fractions were subjected to Glutathione Sepharose 4B (Cyvita, 17-0756-01) for 1.5 h at 4°C. After being washed with cold PBS 3x, Sepharose beads were incubated with PreScission protease (Cyvita, 27084301) in the solution at 4°C to release MAK from the GST tag. Digested products were pelleted, and supernatant fractions were used for *in vitro* kinase assays.

Each reaction for *in vitro* kinase assays is composed of 50 mM Tris-HCl (pH 7.0), 10 mM MgCl_2_, 5 mM ATP (Thermo Scientific, R0441), 1 mM DTT (Thermo Scientific, 20290), 5 µg dephosphorylated myelin basic protein (Merck, 13-110), and purified MAK protein. Reactions were incubated 1 h at 30°C. Samples were mixed with SDS sample dye and incubated for 20 min at 75°C to stop the reactions. Samples were separated by SDS-PAGE and transferred onto PVDF membranes for later analyses. Membranes were blocked with 5% skim milk for 1 h at room temperature followed by probing with primary antibodies and HRP-conjugated secondary antibodies. ECL substrate (Thermo Scientific, 34577) was applied to visualize signals using a iBright FL1500 imager (Invitrogen).

### Data quantification and statistical analysis

To qualify the thickness of photoreceptor layers, photoreceptor layers in the images were straightened with the straighten tool of ImageJ (Fig. S2A). A line dividing the photoreceptor layer into 10 parts was generated with Adobe Illustrator. Photoreceptor layer thickness at each decile was measured with ImageJ by following the conversion ratio of pixels to the actual length of the scale bar. To qualify photoreceptor numbers, photoreceptor layers in the images were straightened with ImageJ, and a reference line marking every 1/10 points was drawn with Adobe Illustrator. Cell numbers were manually counted. TUNEL signals in retinas were manually counted. The zpr1-positive area in the total retinal was calculated as previously described (Fig. S2B) (Nishiwaki et al., 2013). All statistical analyses were performed with GraphPad Prism 9.2.1. Significance levels, numbers of samples, and details of analyses are indicated in each figure and figure legend.

## Acknowledgments

We are grateful to Herwig Baier and Akira Muto for providing the *pday* mutant line. We also thank Tadashi Yamamoto for providing the vector, pGEX6p3. We thank previous lab members, Brandy Lee Denis and Maria Iribarne for supporting the experiment to clone *pday* mutant gene, Shohei Suzuki for cloning of the zebrafish *rhodopsin* promoter, Sachihiro Suzuki for cloning of the zebrafish *gnat2* promoter, Yutaka Kojima, Ayano Harata and Hiroshi Izumi for supporting histological analyses of *pday* mutant phenotypes. We are grateful to the Imaging Section and the sequencing section of the Research Support Division of OIST. We thank Steven D. Aird for critical reading and editing of the manuscript.

## Data availability

N/A

## Author contributions statement

Conceptualization: H-JC, IM; Methodology: H-JC, YN, IM; Software: H-JC, IM; Validation: H-JC, YN, W-CC, IM; Formal analysis: H-JC, YN, IM; Investigation: H-JC, YN, W-CC, IM; Resources: H-JC, YN, W-CC, IM; Data curation: H-JC, YN, W-CC, IM; Writing - original draft: H-JC, IM; Writing - review & editing: H-JC, IM; Visualization: H-JC, YN, W-CC, IM; Supervision: IM; Project administration: H-JC, YN, IM; Funding acquisition: IM.

## Appendix

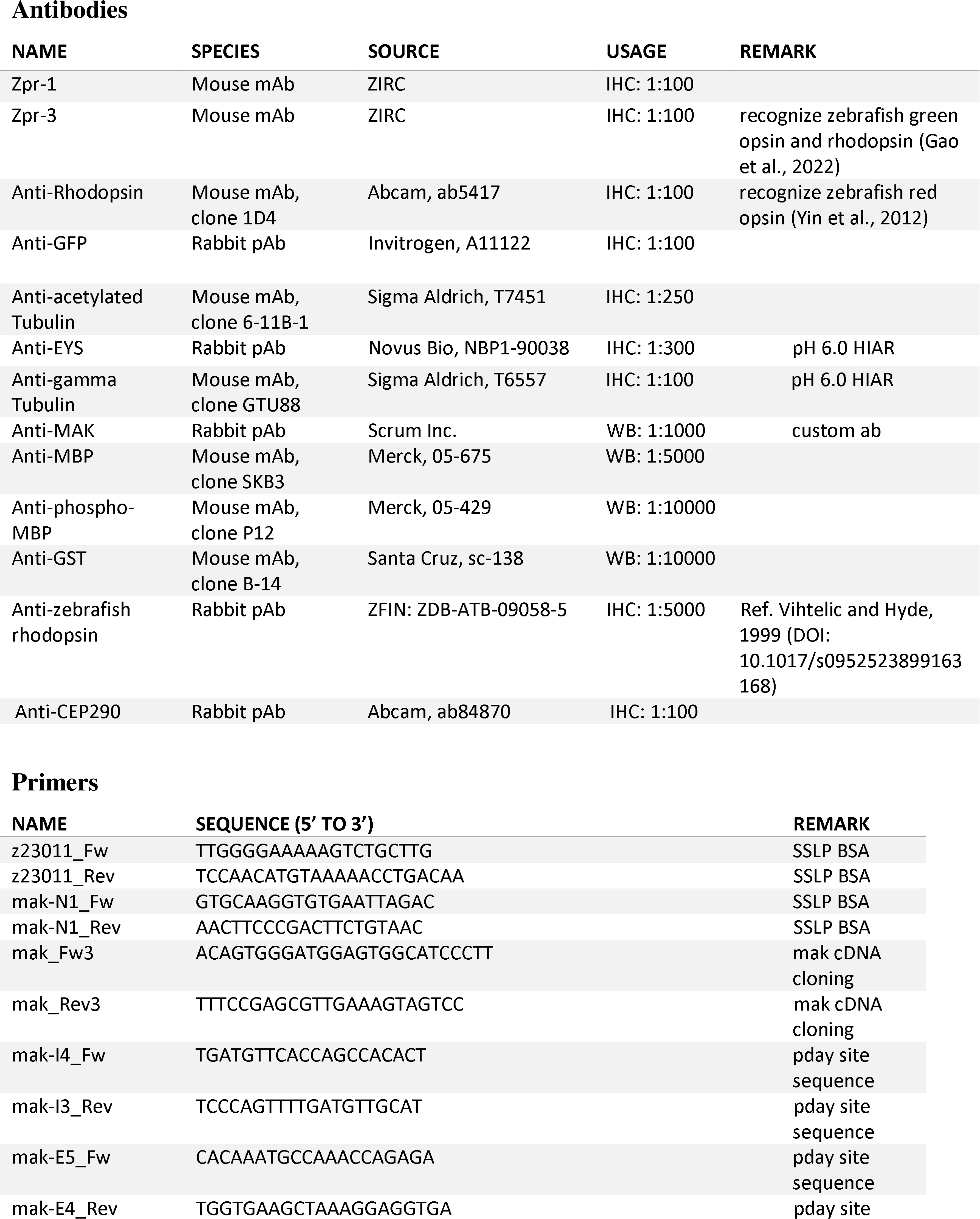
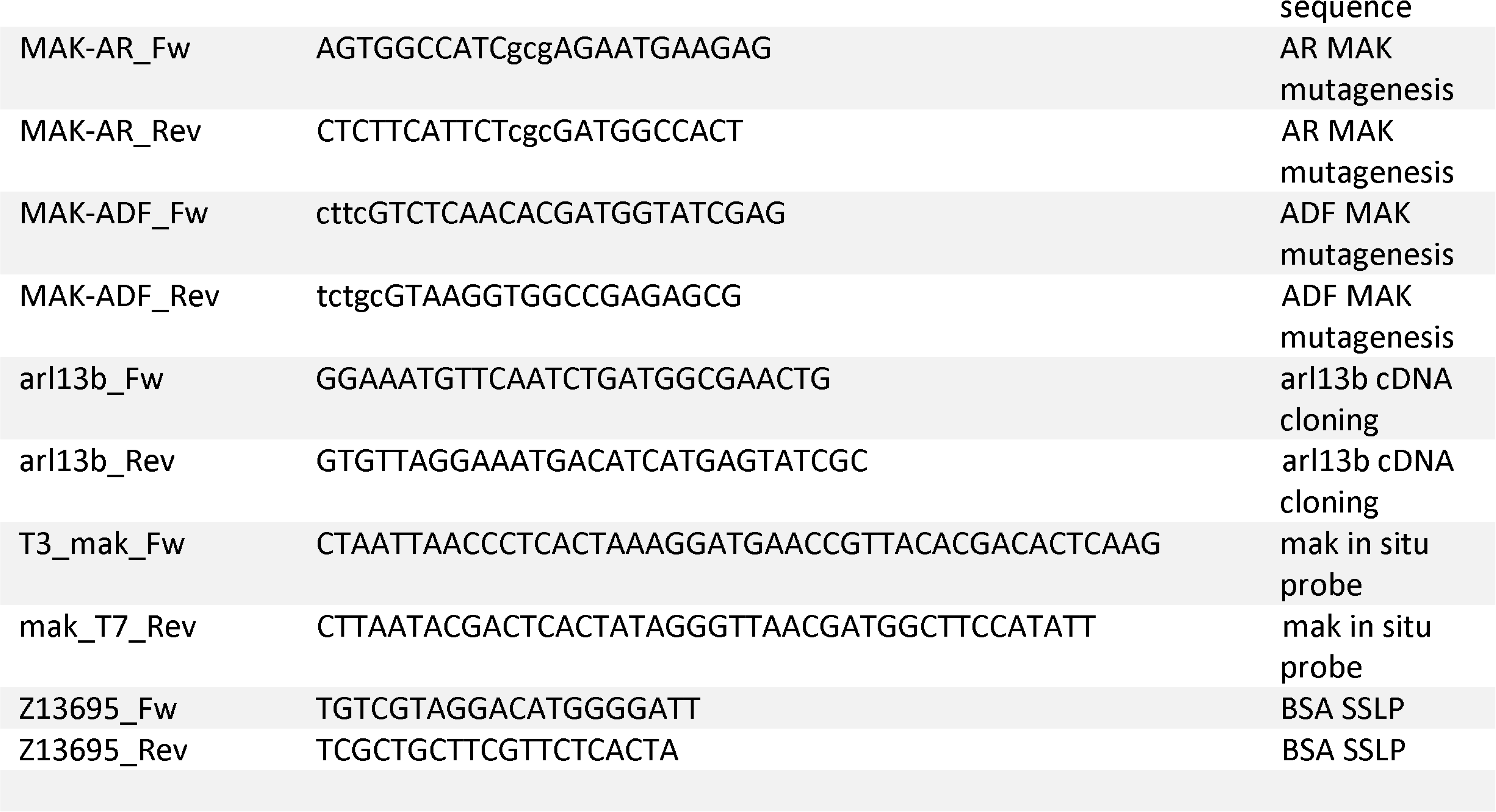

