## Supplementary figures and legends for "Male germ cell-associated kinase (MAK) is required for axoneme formation during ciliogenesis in zebrafish photoreceptors"

**The PDF includes:**
Supplementary figures: Figs. S1 to S5

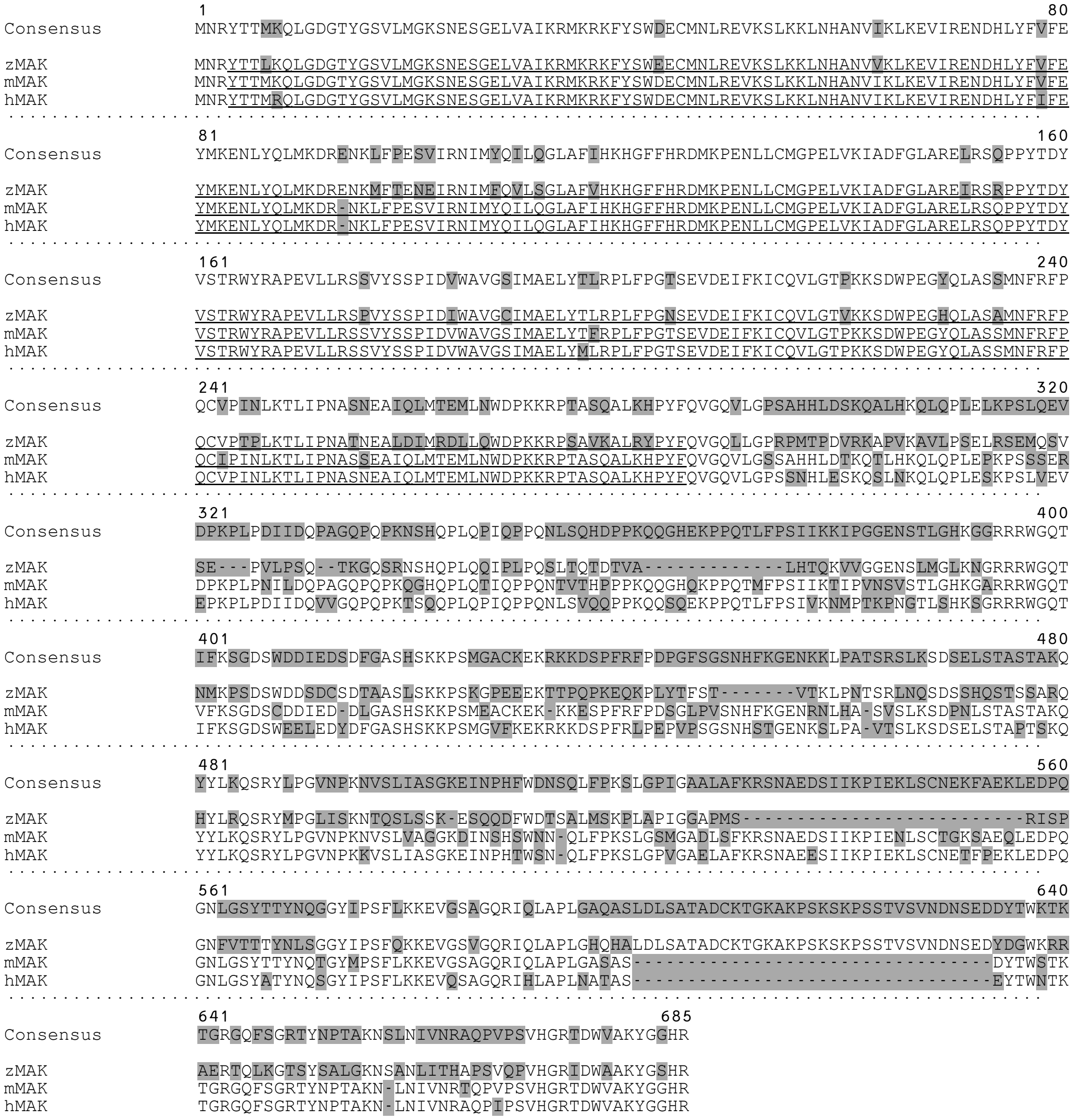

Amino acid sequence identity percentage

| | Full length | Kinase domain | C'-terminus |
|---|---|---|---|
| zMAK vs mMAK | 50% (323/646) | 86.17% (243/282) | 21.82% (79/362) |
| zMAK vs hMAK | 51.38% (333/648) | 88.65% (250/282) | 21.97% (80/364) |
| mMAK vs hMAK | 83.64% (542/648) | 97.86% (275/281) | 72.52% (264/364) |
| Among the three animals | 47.22% (306/648) | 85.10% (240/282) | 17.03% (62/364) |

**Fig. S1: Amino acid comparison between zebrafish, mouse, and human MAK.**
Amino acid sequences of zebrafish, mouse, and human MAK proteins are aligned using Clustal Omega. Grey shading indicates sequence differences. The kinase domain is indicated by an underline. The bottom table shows the percentage of amino acid identity in the full-length protein, the kinase domain, and the C-terminal domain.

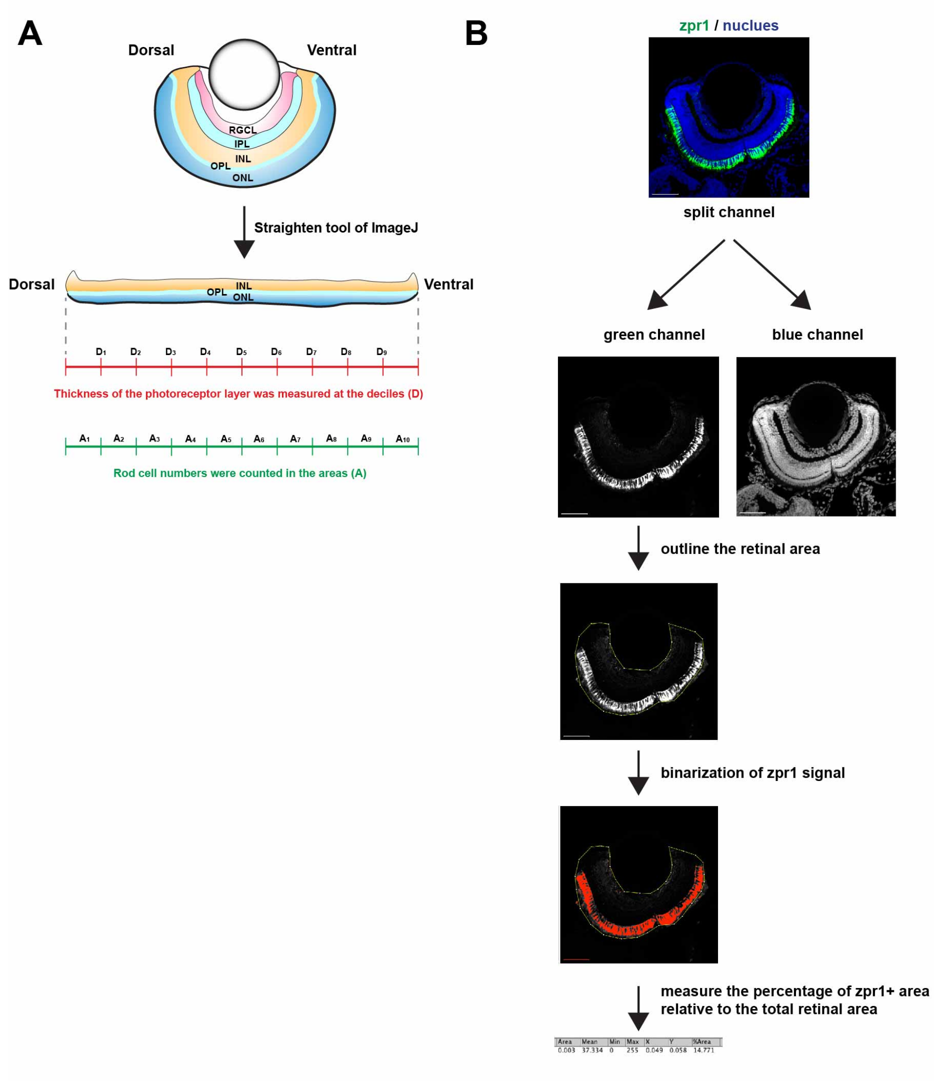


**Fig. S2: Measurement of ONL thickness, rod cell number, and the fraction of zpr1-positive area with regard to total retinal area.**

(A) Images with retinas were straightened with Image J (Fiji). Straightened images were divided into 10 equal areas (A) with 9 deciles (D) between adjacent areas. The thickness of the ONL was measured at the deciles and cell numbers were counted in each area.

(B) Cryosections labeled with zpr1 antibody were scanned under a laser-scanning microscope (LSM510; Carl Zeiss, FV3000; Olympus). First, a one-section image containing the central retina per eye was selected from each individual sample. Using the "split channel" command of Image J (Fiji), confocal scanning RGB color images were split into either R, G or B channel. Only the zpr1 channel (green channel in this figure) is used to draw the outline of the total retinal area. Next, using the "threshold" command of Image J (Fiji), we converted zpr1-positive and -negative areas to a binary scale, 1 and 0, respectively. The number of pixels corresponding to 1 or 0 within the neural retina was determined. Using the "measurement" command of Image J (Fiji), the percentage of zpr1-positive areas relative to the total retinal area was calculated as the percentage of the number of 1 pixels relative to the number of 0+1 pixels. Means and standard deviations were calculated from data obtained for 2-5 sections of more than two embryos.

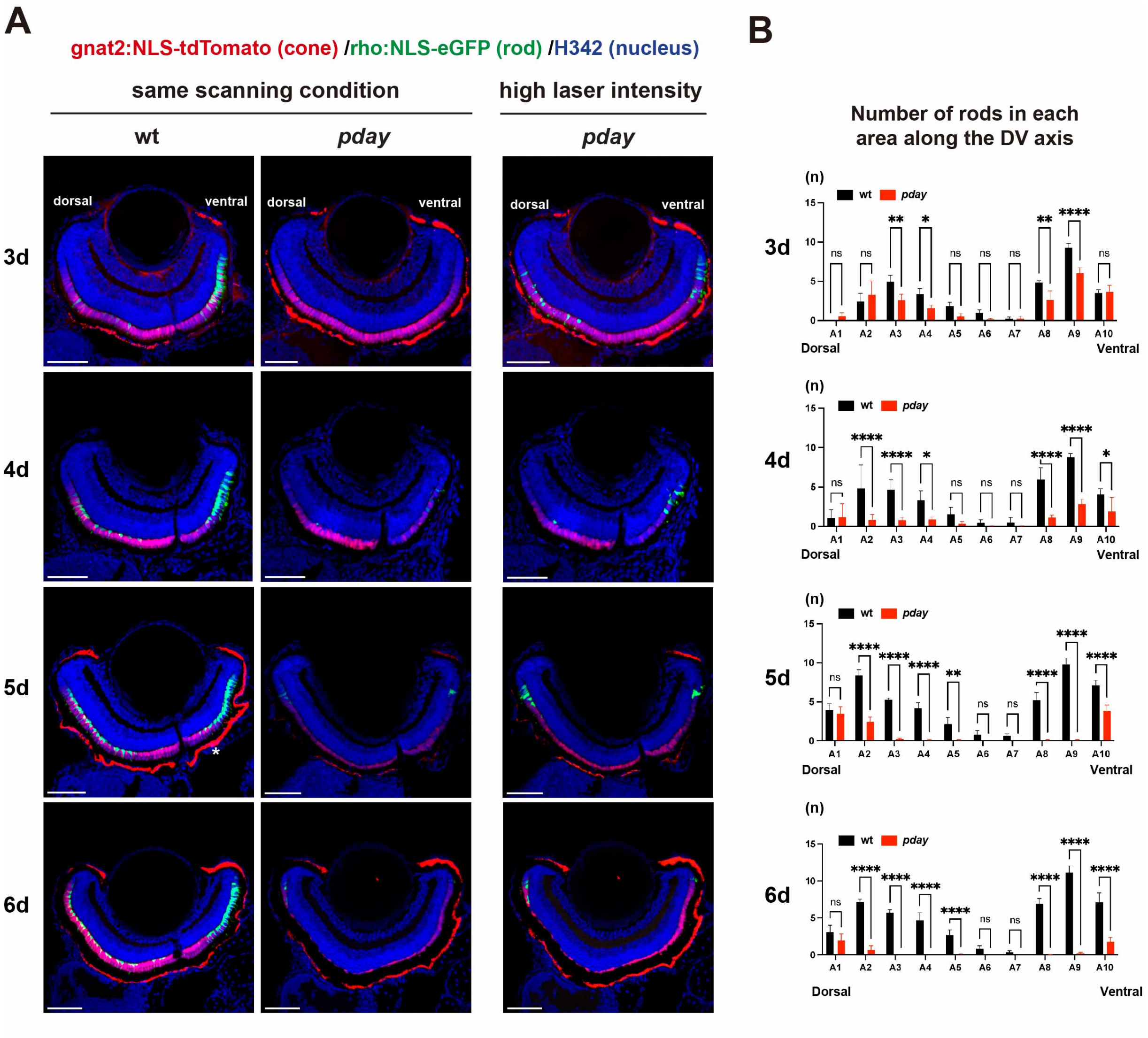


**Fig. S3: Visualization of rods and cones in *pday* mutants.**

(A) Confocal images of wild-type (the left column) and *pday* mutant retinas (middle column) carrying the transgenes *Tg[rho:NLS-eGFP]* and *Tg[gnat2:NLS-tdTomato]*, which visualize rods (green) and cones (red), respectively. Nuclei were counterstained with Hoechst 33342 (blue). In *pday* mutant retinas, eGFP signals were weak. The right column indicates confocal scanning images of *pday* mutant retinas carrying the transgenes *Tg[rho:NLS-eGFP]* and *Tg[gnat2:NLS-tdTomato]*, with higher laser intensity, to enhance eGFP signals. We compare wild-type and *pday* mutant retinas in conventional scanning conditions and with higher laser intensity, respectively, in Figure 2. Scale bars: 40 $\mu$m.

(B) Histogram of rod numbers of wild-type siblings (black) and *pday* mutants (red) retinas at 3, 4, 5, and 6 dpf. Rod number was counted in 1/10$^{th}$ divided area of the ONL along the DV axis and analyzed with 2-way ANOVA and Sidak's multiple comparison tests, n=3 for wild-type, n=4 for *pday* mutants at 3 dpf; n=4 for wild-type, n=5 for *pday* mutants at 4 dpf; n=3 for wild-type, n=3 for *pday* mutants at 5 dpf; n=4 for wild-type, n=4 for *pday* mutants at 6 dpf. Bars and lines indicate mean ± SD. * p<0.02, ** p<0.003, **** p<0.0001, ns: not significant.

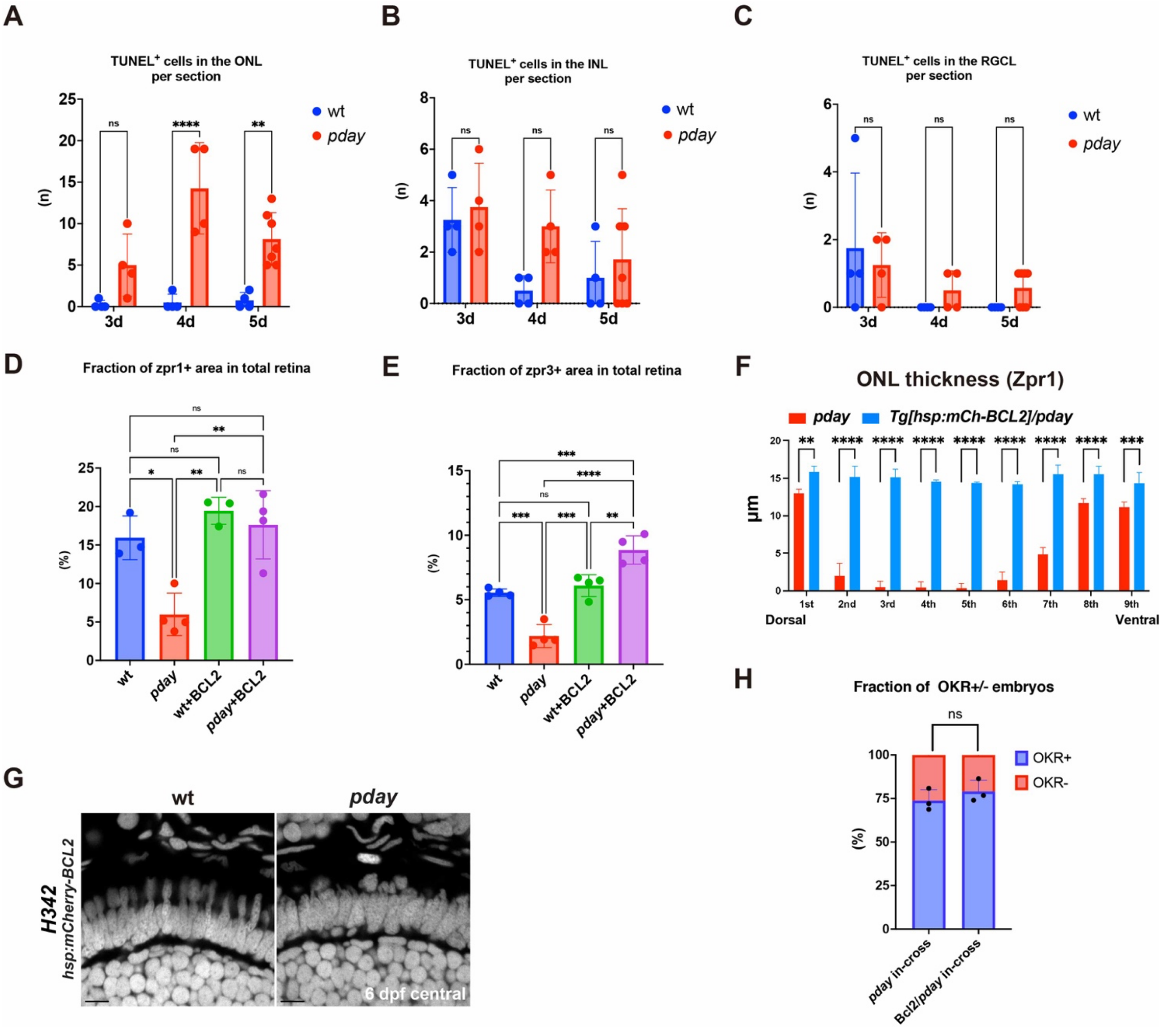


**Fig. S4: Statistical analysis of TUNEL and Bcl2-mediated rescue of photoreceptor phenotypes in *pday* mutants.**

(A) The number of TUNEL signals in wild-type and *pday* mutant ONL at 3, 4, and 5 dpf. TUNEL signals in the ONL are higher in *pday* mutants than in wild type at 3 dpf, although the difference is not significant. TUNEL signals in the ONL are the highest in *pday* mutants at 4 dpf and are still significantly higher in *pday* mutants at 5 dpf, compared with wild-type siblings. Bars and lines indicate mean ± SD. Statistical significance was evaluated with 2-way ANOVA and Sidak's multiple comparison tests; * $p<0.03$, **** $p<0.0001$, ns: not significant.

(B) The number of TUNEL signals in wild-type and *pday* mutant INL at 3, 4, and 5 dpf. TUNEL signals in the INL are higher in *pday* mutants than in wild type at 4 dpf, although the difference is not significant. Bars and lines indicate mean ± SD. Statistical significance was evaluated with 2-way ANOVA and Sidak's multiple comparison tests; ns: not significant.

(C) The number of TUNEL signals in wild-type and *pday* mutant RGCL at 3, 4, and 5 dpf. There is no significant difference between wild-type and *pday* mutants at any stage. Bars and lines indicate mean ± SD. Statistical significance was evaluated with 2-way ANOVA and Sidak's multiple comparison tests; ns: not significant.

(D) The fraction of zpr1-positive area in the total retinal in wild-type and *pday* mutants without and with mCherry-Bcl2 overexpression. The fraction of the zpr1-positive area is markedly reduced in *pday* mutants (red bar), compared with wild-type siblings (blue bar). However, the fraction of the zpr1-positive area is recovered in *pday* mutants with mCherry-Bcl2 overexpression (purple bar), equivalent to wild-type without (blue bar) and with mCherry-Bcl2 overexpression (green bar). Bars and lines indicate mean ± SD. Statistical significance was evaluated with 2-way ANOVA and Turkey's multiple comparison tests; *p<0.0332, **p<0.0021, ns: not significant.

(E) The fraction of the zpr3-positive area in the total retinal area in wild-type and *pday* mutants without and with mCherry-Bcl2 overexpression. The fraction of the zpr3-positive area is markedly reduced in *pday* mutants (red bar), compared with wild-type siblings (blue bar). However, the fraction of the zpr3-positive area is recovered and increased in *pday* mutants with mCherry-Bcl2 overexpression (purple bar), compared with wild-type without (blue bar) and with mCherry-Bcl2 overexpression (green bar), probably due to ectopic distribution of zpr3 signals outside the OS. Bars and lines indicate mean ± SD. Statistical significance was evaluated with 2-way ANOVA and Turkey's multiple comparison tests; **p<0.0021, ***p< 0.0002, ****p<0.0001, ns: not significant.

(F) Histogram of zpr1-positive cone thickness in *pday* mutant retinas with (blue) and without (red) overexpression of mCherry-Bcl2. The ONL is divided into 10 areas along the DV axis. zpr1-positive cone thickness was measured at the interface between each neighboring region and evaluated with 2-way ANOVA and Sidak's multiple comparison tests, n=3 for *pday* mutants, n=4 for *pday* mutants with *Tg[hsp:mCherry-Bcl2]*. ** p<0.003, *** p<0.001, **** p<0.0001. Bars and lines indicate mean ± SD.

(G) Nuclear labeling of wild-type and *pday* mutant ONL overexpressing mCherry-Bcl2 with Hoechst 33342. Nuclear shape in the ONL is slightly irregular in *pday* mutants overexpressing Bcl2.

(H) The percentage of OKR+ (blue) and OKR- (red) embryos produced by crosses of *pday* heterozygous male and female fish without and with the transgene *Tg[hsp:mCherry-Bcl2]*. There is no significant difference between the absence and the presence of *Tg[hsp:mCherry-Bcl2]*. Statistical difference as evaluated with 2-way ANOVA and Sidak's multiple comparison tests. Bars and lines indicate mean ± SD, ns: not significant.

Scale bars: 5 $\mu$m (G).

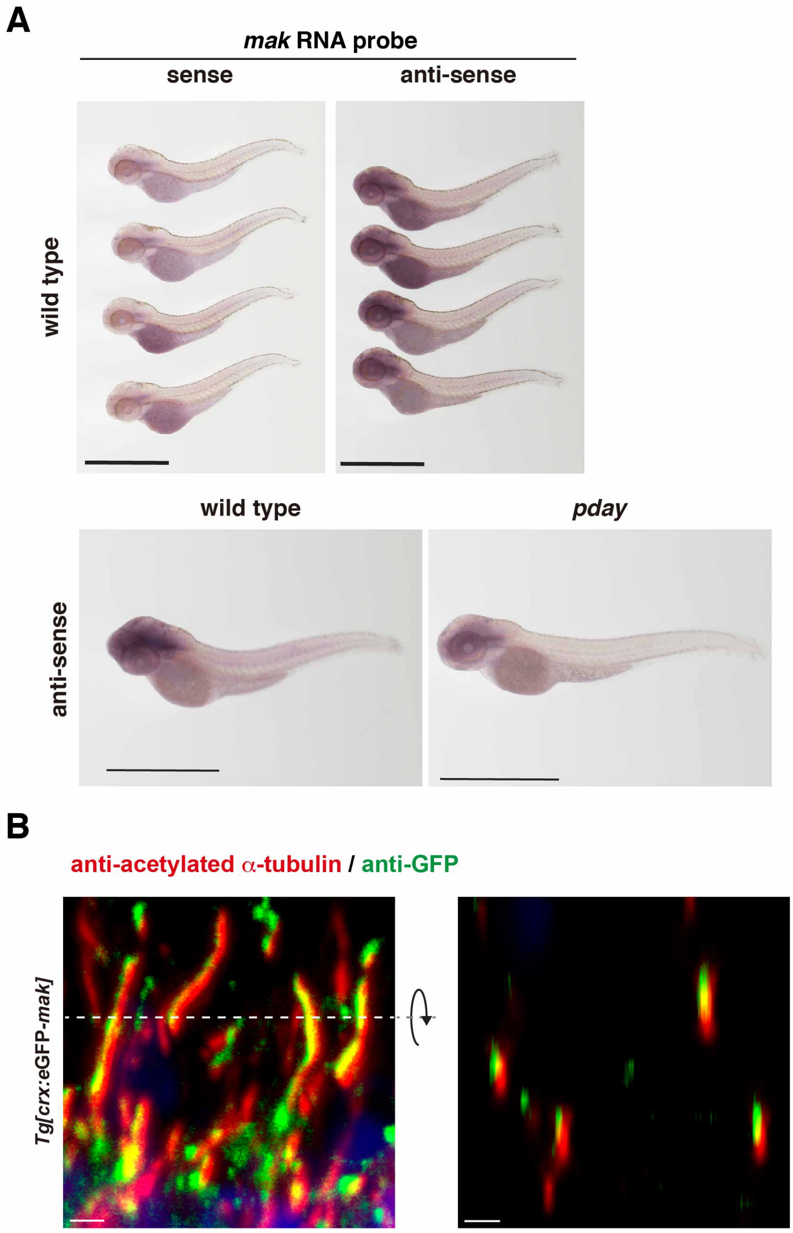


**Fig. S5: *mak* mRNA expression is reduced in *pday* mutants and MAK protein is localized in ciliary axonemes.**

(A) *In situ* hybridization of zebrafish embryos at 72 hpf with *mak* RNA probe. Upper panels indicate *in situ* hybridization with sense (left) and anti-sense RNA probes (right). Bottom panels indicate *in situ* hybridization of wild-type and *pday* mutant embryos with *mak* anti-sense probe. In pday mutant embryos, *mak* mRNA expression is reduced. Scale bars: 1 mm.

(B) Projection view of 3D confocal image of wild-type transgenic *Tg[crx: eGFP-MAK]* ONL with anti-GFP and anti-acetylated α-tubulin antibodies. Nuclei were counterstained with Hoechst 33342. Left panel indicates a radially sectioned projection view of the retina. The eGFP-MAK signals overlap with anti-acetylated α-tubulin signals. Right panel indicates tangentially sectioned projection view, which was digitally produced by rotating the radially sectioned view using Imaris software (Bitplane). Scale bars: 1 μm.
